# Spatio-temporal dynamics of attacks around deaths of wolves: A statistical assessment of lethal control efficiency in France

**DOI:** 10.1101/2024.07.18.604079

**Authors:** Oksana Grente, Thomas Opitz, Christophe Duchamp, Nolwenn Drouet-Hoguet, Simon Chamaillé-Jammes, Olivier Gimenez

## Abstract

The lethal control of large carnivores is criticized regarding its efficiency to prevent hotspots of attacks on livestock. Previous studies, mainly focused on North America, provided mixed results. We evaluated the effects of wolf lethal removals on the distribution of attack intensities in the French Alps between 2011 and 2020, using a Before After Control-Impact approach with retrospective data. We built an original framework combining both continuous spatial and temporal scales and a 3D kernel estimation. We compared the attack intensities observed before and after the legal killings of wolves on a period of 90 days and on a range of 10 km, and with control situations where no removal occurred. The analysis was corrected for the presence of livestock. A moderate decrease of attack intensity was the most common outcome after the lethal removal of a single wolf. This reduction was greatly amplified when removing two or three wolves. The scale of analysis also modulated this general pattern, with decreases being generally amplified at a small spatio-temporal range. Contextual factors (*e.g.*, geographical or seasonal variations) could also lead to deviations from this general pattern. Overall, between 2011 and 2020, lethal control of wolves in France generally contributed to reduce livestock attacks, but mainly locally and to a minor extent. Our results highlight the importance of accounting for scale in such assessments and suggest that the evaluation of the effectiveness of lethal removals in reducing livestock predation might be more relevant in a local context. As a next step, we recommend to move forward from patterns to mechanisms by linking the effects of lethal control on wolves to their effects on attacks through analysis of fine-scaled data on wolves and livestock.

## 1. Introduction

Conservation of large carnivores while limiting their damages on livestock is a difficult task worldwide (van Eeden et al., 2018). Predation on livestock by large carnivores has been the main fuel of hostility towards them (Lute et al., 2018; van Eeden et al., 2018). Strategies of eradication led to their local extirpations in some cases (Treves and Karanth, 2003), like in many areas across Europe (Chapron et al., 2014). The growing promotion of biodiversity conservation these last decades has reversed the trend, with some populations of large carnivores now expanding in Europe, though at unequal paces (Kaczensky et al., 2024). Diverse measures have been promoted and implemented with the objective of mitigating or limiting predations, whether by increasing acceptance through compensations (Gervasi et al., 2021), by decreasing livestock vulnerability through changes in husbandry practices, or by directly affecting carnivores (van Eeden et al., 2018).

Lethal control, one of the most controversial measures targeting carnivores, aims to decrease carnivore density and so the encounter risk with livestock, or to remove specific individuals inclined to predation on livestock, while maintaining their conservation status as favorable (Lennox et al., 2018). Lethal control has been widely applied worldwide, and can be exclusively directed towards carnivores attacking livestock or be part of a more general harvest (Lorand et al., 2022; Treves and Karanth, 2003).

The side effects of lethal control are often claimed to undermine its efficiency or even to create more damages (Elbroch and Treves, 2023). In the specific case of social canids like wolves, studies have mostly pointed out additive human-caused breeder loss as a risk for increasing damages on livestock (Jęodrzejewska et al., 1996; Žunna et al., 2023). Social instability can indeed cause multiple reproduction (Ausband et al., 2017b) or pack disruption (Cassidy et al., 2023), although their effects on damages have not been assessed.

Implementing randomized controlled experiments to evaluate the efficiency of lethal control of large carnivores is extremely difficult, considering the involved large spatial and temporal scales. To date, no experimental study has been conducted for this matter (Treves et al., 2016, 2019). Only Allen (2014) conducted a controlled experiment to evaluate dingo’s baiting efficiency but using the number of lactation failures as a proxy for calves lost to predation instead of direct cases of predation.

Also, retrospective studies have been conducted to answer the question of lethal control efficiency. The data they used were not collected within a scientific frame but generally through management (*e.g.* damages from compensation schemes). Most of retrospective studies on social canids (gray wolves *Canis lupus*, dingoes *Dingo dingo*, coyotes *Canis latrans*) were correlative and tested for a relationship between the number of livestock damages and the number of killed predators (Šuba et al., 2023; Fernández-Gil et al., 2016; DeCesare et al., 2018; Conner et al., 1998; Allen, 2015; Kutal et al., 2023; Merz et al., 2025). However, the ability of this approach to infer causality is being debated (Arif and MacNeil, 2022; Nichols and Cooch, 2025). The approach is also subject to analytical variability (Gould et al., 2025). For example, three correlative studies which used the same dataset concluded differently on the effect of lethal removals of wolves on livestock damages (Kompaniyets and Evans, 2017; Poudyal et al., 2016; Wielgus and Peebles, 2014).

A smaller number of retrospective studies used a comparative approach. Harper et al. (2008), Bradley et al. (2015), Santiago-Avila et al. (2018) and Wagner and Conover (1999) compared situations where lethal control was used and situations where it was not (*i.e.* inter-site comparison). Bjorge and Gunson (1985) and Blejwas et al. (2002) compared the situation before and after the implementation of lethal control within the same site (*i.e.* intra-site comparison). While the inter-site comparison is more susceptible to possible spatial confounding factors but minimizes temporal confounding factors, the opposite is true for the intra-site comparison. Kutal et al. (2023) was the only study to mix inter-site and intra-site comparisons, by combining comparisons before and after hunting season and between sites with and without hunted wolves (*i.e.* Before-After Control-Impact, BACI). The BACI study design showed a better performance to estimate the effect of an impact in a simulated ecological dataset than intra-site or inter-site comparisons (Christie et al., 2019), and performed even better than the randomized controlled experiment still considered as gold-standard of scientific inference (Treves et al., 2019).

Five of these seven comparative studies on social canids showed that lethal control significantly decreased damage recurrence, whereas Santiago-Avila et al. (2018) and Kutal et al. (2023) found no significant effect of lethal control on damages. Yet, results suggested variability in the efficiency of lethal control. For example, Harper et al. (2008) detected that only the death of an adult wolf male was significantly efficient, whereas Bradley et al. (2015) did not detect sex or age conditions on efficiency. Variability could also result from the study designs themselves, as these studies used very different spatial and temporal scales to evaluate lethal control efficiency. Thus, small scale (*e.g.* 4 km in Harper et al. (2008)), medium scale (*e.g.* pack territory in Bradley et al. (2015)) or large scale (*e.g.* 711 km² in Kutal et al. (2023)) could be used for space, sometimes within the same study (*e.g*.

Santiago-Avila et al., 2018). Time scale could also be disparate, going from several days or months (*e.g.* Blejwas et al. (2002)) to one year (*e.g.* Kutal et al., (2023)) or more (*e.g.* Bjorge and Gunson (1985)). Most importantly, none of the mentioned retrospective studies have both used continuous spatial and temporal scales, making it difficult to identify potential gradients of lethal control effects, as those suggested in Santiago-Avila et al. (2018). Finally, only two of the seven comparative studies have controlled for livestock presence or abundance (Blejwas et al., 2002; Wagner and Conover, 1999), yet it is the primary factor structuring damage risk and the highest risk of confounding factor.

The aim of our study was to evaluate the effects of the lethal control applied for gray wolves in France on the intensity of attacks on sheep. We used the data collected by the French administrations for the period 2011-2020. Given the retrospective nature of our study, we attempted to fill the gaps identified in the previously mentioned retrospective studies on the subject. For that purpose, we used the study design of BACI, used continuous spatial and temporal study scales and corrected for livestock presence through an original framework of kernel estimation.

## 2. Methods

### 2.1 Management context and study area

After historical campaigns of destruction, gray wolves disappeared from France at the beginning of the twentieth century, despite being present on the whole metropolitan territory at the end of the eighteenth century (de Beaufort, 1987). From the 1970s, the combination of protective legislation and the increase of forested habitat allowed the recovery of the species in Europe (Chapron et al., 2014). In the 1990s, wolves from the Apennines population in Central Italy dispersed and settled into the south-western Alps, straddling Italy and France (Linnell and Cretois, 2018; Fabbri et al., 2007; Poulle et al., 1997). From there, a wolf population progressively established in the area. On the French side, the population has expanded but has until now remained mainly restricted to the south-eastern corner of France, except for a few reproductive packs outside of this area (Louvrier et al., 2018). The spatial growth reflected a demographic growth that has been exponential. In the first twenty years of the recovery, the population size progressively increased and reached approximately 200 estimated individuals in 2013 (Réseau Loup-Lynx, 2024). Then, the growth rate accelerated, and the population more than doubled in five years, with more than 500 estimated individuals in 2018 (Office National de la Chasse et de la Faune Sauvage, 2019). This sustained growth has continued at least up to 2022, with a population size approaching 800 estimated individuals (Milleret et al., 2025a).

South-eastern France is characterized by a variety of landscapes, including high altitude mountains, Piedmont plains, hills and garrigue lands, where mountain and Mediterranean climates converge. Livestock farming is generally extensive. Herds of cow are principally present in the north of the region, while those of sheep are present in the whole area (Dobremez et al., 2016). The pastoral practice of transhumance, consisting in moving all or part of herds from low to high altitude pastures during summer and vice versa during winter, is still important. Duration of grazing season increases following a north-south gradient, with some herds from low altitudes in the south grazing outside all year round (Dobremez et al., 2016). Livestock owners can share pastures, and even gather their herds to limit the cost of transport for transhumance and of herding. In the Alps, herds of cows are generally not mixed with flocks of sheep, but the latter can be occasionally mixed with goats. Mean size of transhumant flocks of sheep varies greatly between alpine departments and ranges between less than 250 sheep up to 2000 sheep (Agreste, 2021).

To prevent attacks, the authorities subsidize non-lethal measures for protecting flocks of sheep or goat through livestock guarding dogs, fences and additional shepherds. In situations where attacks persist despite non-lethal measures, the authorities have authorized lethal removals of wolves by ground shooting since 2004, based on the article 12 of the Directive Habitats. Other forms of removals (*e.g.* trapping, aerial shooting) are prohibited. Conditions of application of ground shooting are defined by ministerial decrees (*e.g.* Ministère de la Transition Ecologique et Solidaire, 2018) and depend on the implementation of non-lethal protective measures and on the recurrence of attacks. If the conditions are met, the administration assigns the derogation to the livestock owner for a duration between two months and five years. The owner itself or a mandated hunter can shoot wolves during an attack on its livestock (hereafter ‘simple defense’). When attacks still remain, governmental agents or special volunteers who are trained may support owners in protecting their livestock (hereafter ‘reinforced defense’). Eventually, if the level of attacks escalates in an area, the administration can decide to shoot a specific number of wolves under game hunting conditions (hereafter ‘hunting’). Between 2004 and 2018, the number of wolves that could be annually killed if required was prevented from exceeding between 10 and 12% of the annual estimated winter wolf population size, in order to avoid threatening the recovery of the species. Throughout this period, the number of killed wolves followed the increases of both the wolf population size and the attack occurrences, and from 2011, wolves have been legally killed every year (Grente, 2021). Since 2019, considering that the wolf population has reached a satisfying demographic conservation status, France has increased the possible number of lethal removals to 21% of the annual winter population size estimate (Ministère de la Transition Ecologique et Solidaire, 2020).

Because wolves were removed sporadically between 2004 and 2010 (range: 0-2 wolves per year), we started our study period in 2011, until 2020 included. Within the study period, only three lethal removals were outside south-eastern France and concerned lone wolves, which dispersed far and whose territories were very isolated. Their deaths could only lead to an absence of attacks. This context was totally different from the one in the Alpine range where the wolf population was settled. Therefore, we restricted our study area to the two south-eastern regions, Provence-Alpes-Côte-d’Azur in its entirety and Auvergne-Rhône-Alpes restricted to its five most south-eastern departments, Ardèche, Drôme, Isère, Savoie and Haute-Savoie (Figure 1).

**Figure 1.**
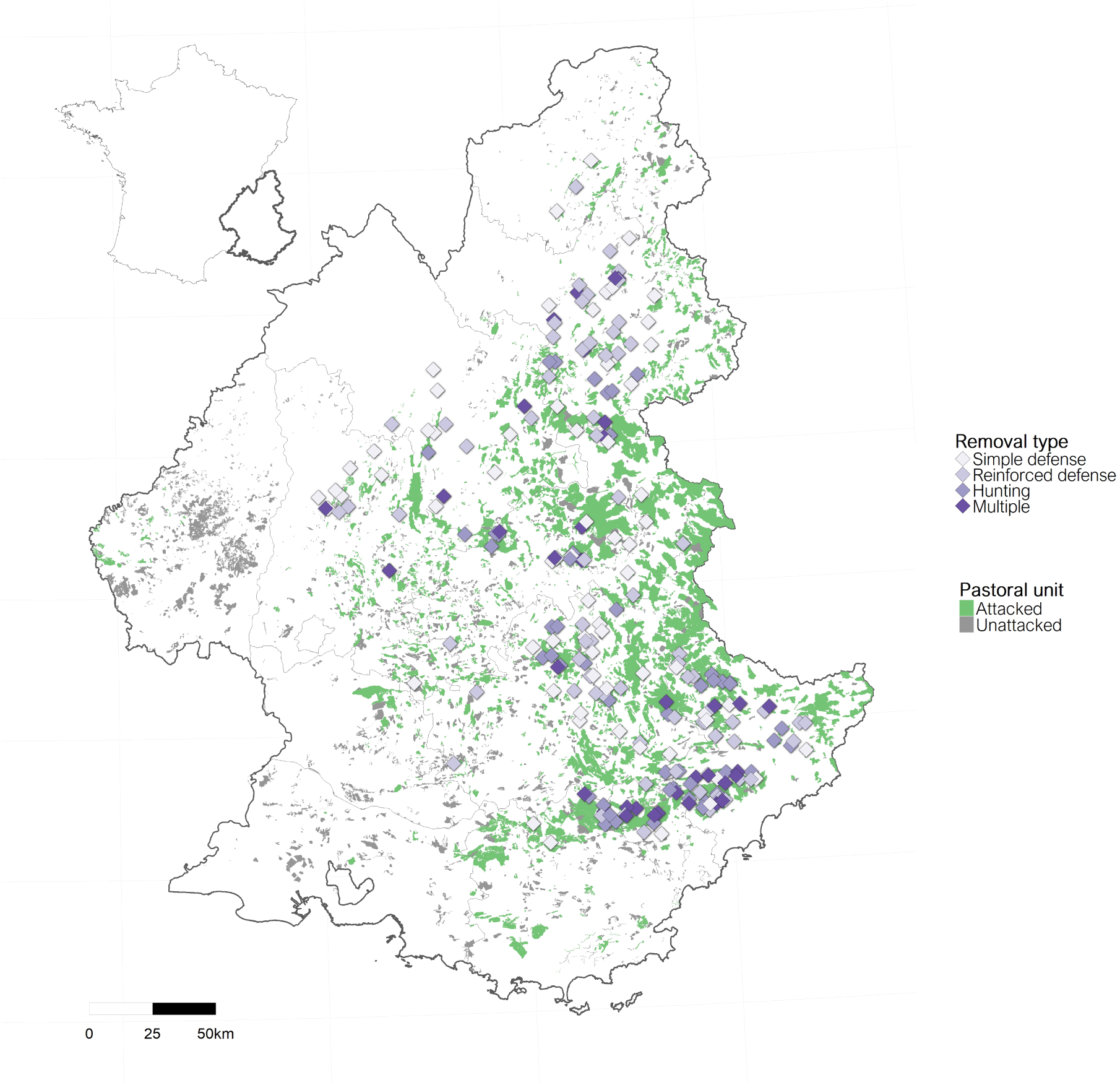
Study area and location within France. Purple points represent locations of the 188 single lethal removals, colored according to the three administrative types of shooting implementation (simple defense, reinforced defense, hunting) and of the centroids of the 35 multiple lethal removals (darkest color). Pastoral units with or without attacks are shown in green and grey, respectively.

### 2.2 Datasets

#### 2.2.1 Lethal removals

We used the 362 administrative records of wolves killed by legal shootings from 2011-2020 that were geolocated within the study area. Dates of lethal removals were recorded, but not the hour. The number of lethal removals increased over time and could be divided into three periods, 2011-2014, 2015-2018 and 2019-2020, that represented respectively 8%, 44% and 48% of the dataset, for annual means of 7 (standard deviation, hereafter ‘±’, 6), 40 (± 6) and 87 (± 3) lethal removals respectively.

Almost half of the records were cases of reinforced defense (48%), while simple defense and hunting cases represented respectively 27% and 25% of the dataset. There were 58 records (16%) for which no wolf body was found after the shooting, but they were considered as killed because there was enough evidence for lethal injuries. For the remaining 304 records, information from the necropsy reports was available on the sex and the age class (pup, subadult or adult) of the killed wolves and on the annual reproductive status for females.

#### 2.2.2 Attacks on sheep

We used the 21 936 claims of attacks on sheep from 2010-2020 and geolocated within the study area, that were compiled by the French administration and for which wolf responsibility could not be discarded. Each claim corresponded to one or several dead or wounded sheep.

Apart very few specific cases, all claims were checked in the field by trained public agents. They followed a standardized procedure to estimate the date of the attack and if wolf responsibility could be discarded or not. Compensation was attributed regardless of non-lethal measures up to 2019. From 2020, compensations after two compensated attacks depended on the implementation of protective measures, with conditions of implementation varying locally according to the level of attacks of municipalities. Protective measures are widely implemented in south-eastern France (Meuret et al., 2020). We have therefore confidence in the level of reports of attacks noticed by breeders.

#### 2.2.3 Spatio-temporal sheep distribution

We used the data from the pastoral census of the two regions Provence-Alpes-Côte-d’Azur and Auvergne-Rhône-Alpes conducted by the National Research Institute for Agriculture, Food and the Environment (Dobremez et al., 2016). The census indexed and delimited for the period 2012-2014 the pastoral units at medium or high altitude used by flocks, as well as the pastoral zones at lower altitudes. We restricted the dataset to units and zones that were used for grazing sheep, regardless of the other livestock. The annual number of grazing sheep was on average 955 (± 764) for units (range 2-6000) and 412 (± 411) for zones (range 1-3600). The months for which the area was grazed were known for each pastoral unit. Only the seasons in use were available for pastoral zones.

Therefore, we attributed to the pastoral zones the months corresponding to the seasons for which they were grazed (spring: March to May, summer: June to August, fall: September to November, winter: December to February). The annual number of grazed months was on average 6.9 (range 1-12). The total size of our final dataset of pastoral units and zones (hereafter ‘pastoral units’) was 4987 (Figure 1).

A substantial part of the attacks (41%) was not occurring within pastoral units during their set of grazed months. This part was only reduced at 35% when considering the attacks from the period of the pastoral census (2012-2014). Therefore, most mismatches were not due to the temporal desynchronization between the two datasets. These mismatches certainly resulted from changes in the use of pastures that shepherds or breeders need to apply to adapt to real-time conditions. We corrected minor mismatches in the attack and pastoral datasets by following the procedure described in Supplementary Information S1. We excluded 13% of the attack dataset from the analysis, because no pastoral unit could be found within 500 meters from these attacks, or because they occurred in units that were supposedly not used, with a temporal mismatch of more than 3 months. The final dataset of attacks had 19 302 events. We discuss the limits of this correction in the Discussion section.

### 2.3 Analysis

Apart from being precisely geolocated, our data about attacks and lethal removals were very unbalanced in favor of attacks (19 302 attacks against 362 lethal removals). Consequently, we adopted an original approach consisting in choosing removals as focal points, and in computing attack intensities over continuous spatial and temporal scales.

#### 2.3.1 Spatio-temporal distributions of attacks around removals

We grouped the lethal removals separated by less than 5 km and less than 25 days (hereafter, ‘multiple removals’), and considered the other removals as independent (hereafter, ‘single removals’).

For each single removal, we only selected the attacks in a radius of 10 km from its location (Figure 2) and across 90 days before and 90 days after its date. The day of the removal was not part of the buffer, because we could not know if an attack recorded at the date of the removal occurred after or before the removal, having no information about the hour of both attacks and removals. We attributed two coordinates *(i, j)* to each attack (Figure 2). Coordinate *i* corresponded to the spatial distance *i* (km) between the attack and the removal. Coordinate *j* corresponded to the temporal distance (days) between the date of the attack and the date of the removal. The point of coordinates *i* = 0 and *j* = 0 represented the spatial location of the removal, and the day before the removal (*i.e.* the day of the removals was not part of the analysis).

**Figure 2.**
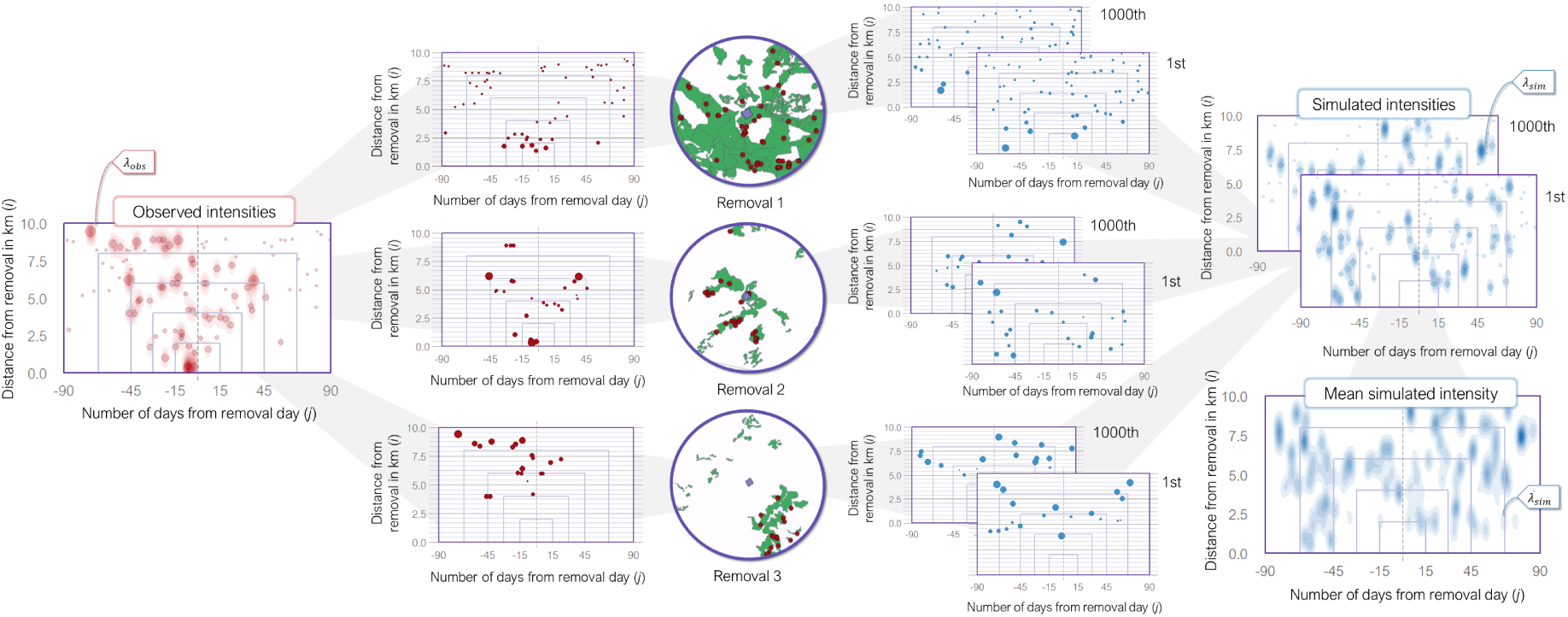
Example of the spatio-temporal distributions of attacks around three lethal removals. The circles in the center show the spatial distribution of observed attacks (red points) within the pastoral units (green polygons) 10 km from the removal location (purple points). The intermediate graphs show the distributions of the observed attacks (left) and of two sets of simulated attacks (right) around removals along a unique spatial axis (i) and temporal axis (j). The size of points represents the spatial correction (see Supplementary Information S2). The observed and simulated attack intensities λ_obs_ and λ_sim_ on the sides are calculated by aggregating the observed attacks and the simulated sets of attacks, respectively. The mean simulated attack intensities λ_sim_ are eventually calculated by averaging λ_sim_(i,j) of all simulated sets (here only two for the example). Rectangles delimit the five nested scales for trend computations: 15, 30, 45, 67 and 90 days and 0, 2, 4, 6, 8 and 10 km.

In the case of multiple removals, we selected the attacks in a radius of 10 km from the centroid of the removals of the group, and across 90 days before the date of the first removal of the group and 90 days after the date of the last removal of the group. As before, we attributed the same two coordinates *(i, j)* to each attack. Coordinate *i* corresponded to the spatial distance *i* (km) between the attack and the centroid of the group. Coordinate *j* corresponded to the temporal distance (days) between the date of the attack and the date of the first (or last) removal of the group if the attack occurred before (or after) the period covered by the group.

Attributing one spatial coordinate *i* to the attack instead of two traditional coordinates induces a spatial bias that needed to be corrected.

We ensured that the spatial and temporal selections of the attacks for each removal did not overlap areas or periods without data about attacks. We excluded from the analysis the 84 lethal removals occurring at less than 10 km from the limits of the study area or occurring after the 1^st^ October 2020.

Eventually, we randomly simulated attacks around each removal, with a distribution defined according to the locations of pastoral units within 10 km and their grazed months. The simulated attack dataset corresponded to the expected distribution of the attacks if they only depended on the livestock presence. We repeated the simulation 1000 times (Figure 2).

More information about the selection of attacks, the correction of the spatial bias and the simulation of attacks is available in Supplementary Information S2.

#### 2.3.2 Intra-site comparison

To generalize the spatio-temporal distributions of the attacks around removals, we aggregated the observed attacks of all removals into a unique set (Figure 2). We repeated the same process for each of the 1000 replications of simulated attacks. We treated separately the dataset of single removals and the dataset of multiple removals. We also explored other sets of aggregation of single removals (hereafter, ‘subsets’) that we described in the next section.

We then estimated the observed and simulated attack intensities *λ_obs_*and *λ_sim_* by applying an anisotropic Gaussian kernel density estimation (Figure 2). Given similar spatial corrections, the higher the number of attacks within the vicinity of the same pair of coordinates (*i,j*), the higher the attack intensity estimate of the related pixel *λ(i,j)*. Eventually, we computed the mean simulated attack intensities *λ*_sim_ by averaging *λ_sim_(i,j)* across the 1000 sets of simulations.

To correct the observed intensities *λ_obs_* for livestock presence, we calculated the corrected intensities *λ_corr_(i, j)* as:

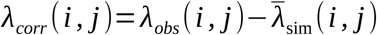

Thus, a value of *λ_corr_(i, j)* lower, equal or higher than zero indicated that the observed attack intensity at coordinates *(i, j)* was lower, equal or higher, respectively, than the attack intensity expected according to pastoral use.

Eventually, we calculated the trends of observed or corrected attack intensities after removals, taking the intensities before removals as the basis. Both observed and corrected trends could exceed 100% if the total of intensities after removals more than doubled the total of intensities before removals. Corrected trends could also be lower than -100% because *λ_corr_* could take negative values. The standard deviation of each trend was estimated by using the jackknife method (Maesono, 2011).

First, we calculated trends at five nested scales, by progressively increasing both spatial and temporal scales (Figure 2). We also calculated these trends by increasing the temporal scale only, for three fixed spatial scales. Second, we investigated the potential spatial shift of attack intensities by calculating trends at each spatial unit for three fixed temporal scales. Contrary to the nested trends, we calculated the trend of the attack intensity at *j* km, not from 0 to *j* km. In all cases, trends were adjusted to allow comparison across scales or sets of removals.

More information about the attack intensity estimation and the calculations of trends is available in Supplementary Information S3.

#### 2.3.3 Inter-site comparison

Lastly, we combined the intra-site comparison performed in the previous section with an inter-site comparison. In other words, we applied a complete BACI study design (Christie et al., 2019). This study design allowed us to address confounding factors but also temporal autocorrelation, *i.e.* the risks that attacks naturally persisted, increased or decreased after lethal removals, without being related to the application of lethal removals. The correction of attack intensities for livestock presence allowed us to partly address the risk of confounding factors, because livestock presence is the primary factor of attack distribution, but there were many other possible factors that could modify attack distribution beside lethal removals (*e.g.* livestock protection, wild prey distribution, wolf poaching…).

We compared the trends of the spatio-temporal distributions of attacks around lethal removals and those around control events where no lethal removal was applied. If the trends of lethal removals and of control events were similar, we could assume that the trends after lethal removals were not the result of the removals themselves, but the result of confounding factors or of temporal autocorrelation. Otherwise, we could assume that the lethal removals mainly caused the observed changes on attack distribution after their application.

Each removal received one control event. In the case of a single removal, we attributed the date of the removal to its control event. In the case of a multiple removal, we attributed two dates to its control event: the date of the first removal of the group and the date of the last removal of the group.

Then, we delimited the sampling area where the spatial location of the control event could be randomly sampled. We considered that only the geographic zone the removal was within represented a similar environment. We delimited these zones by using 25 Alpine mountain ranges defined by hydrography and orography (Knopf, 2019) and the limits from OpenStreetMap of the protected area *Parc Naturel Régional des Préalpes d’Azur* (PNRPA) and of the military zone *Canjuers*. We buffered the zones by 15 km, in order to include removals in their vicinity, provided that the buffer did not overlap another buffered zone, or the area closer than 10 km from limits of the study area (Figure 3). We excluded from the analysis the 9 removals that could not be attributed to a zone. However, the attack distribution close to a control event could be distorted by the lethal removals occurring shortly before or after the control event. Therefore, the sampling area did not include the area within a 10-km radius around any known removal occurring less than 90 days from the control event.

**Figure 3.**
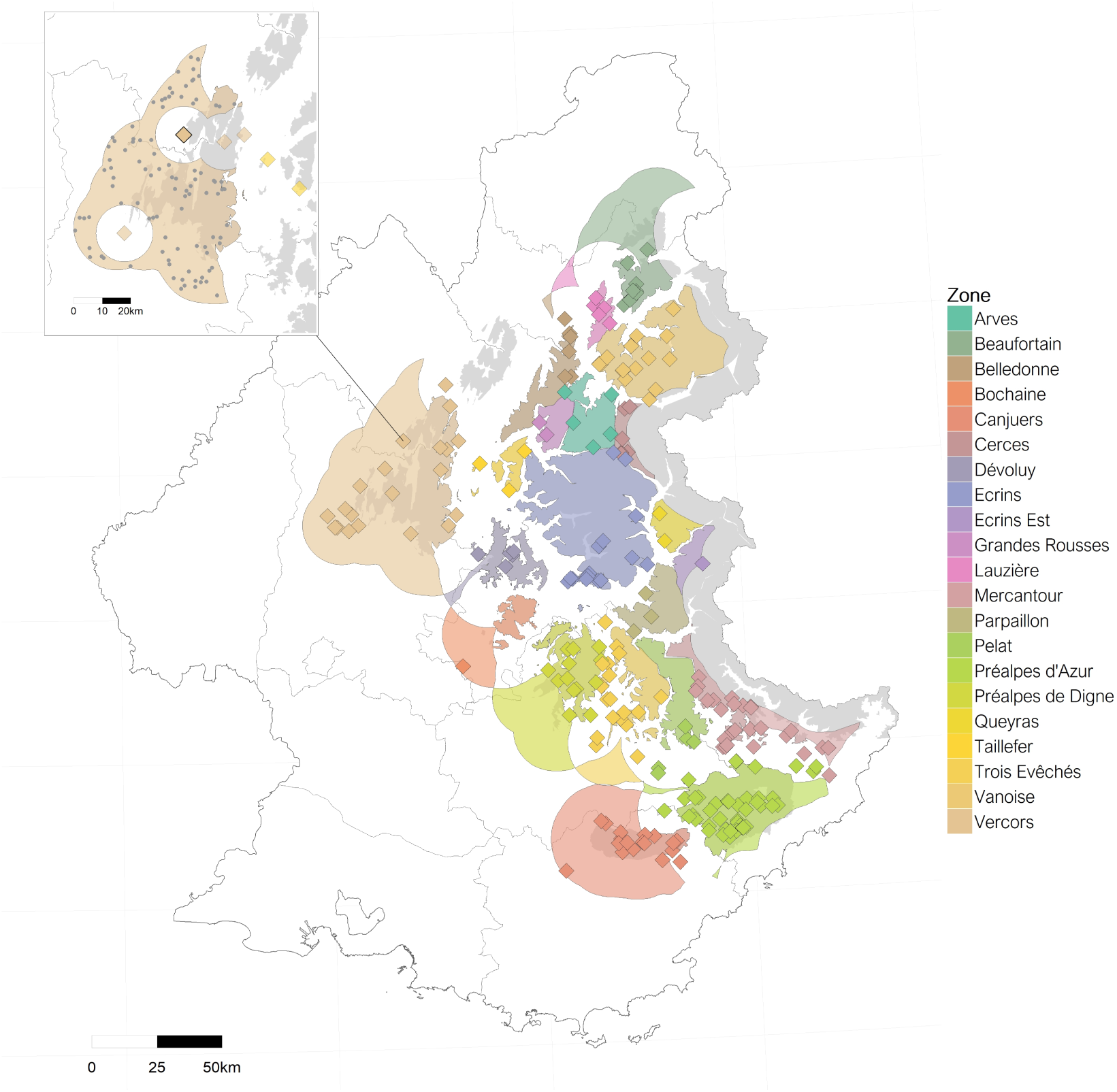
Sampling areas for the control events of the 223 lethal removals (colored squares). The inset plot shows the sampling area of one focal removal (thick square) in the Vercors zone, and its 100 sampled control events (grey round points).

In total, we generated 100 control events per lethal removal. In the same way as for the sets of lethal removals, we applied the intra-site comparison process previously described to each generation of control events (hereafter, ‘control set’).

Eventually, we assessed if the removals significantly modified the intensities of attack intensities (*i.e.* alternative hypothesis), or if there was no sufficient evidence to reflect an effect of removals on attacks (*i.e.* null hypothesis). We considered that the effect was significant if the confidence intervals of the trends following removals were not overlapping the confidence intervals of the trends following control sets. Following Knol et al. (2011), we adapted the levels of the confidence intervals (range 83.4–95%) in order to set the significance level at 0.05 (*i.e.* 5% of risks of rejecting the null hypothesis when it is true).

In the end, we analyzed 188 single removals (1 killed wolf) and 35 multiple removals, with on average 2.29 killed wolves (± 0.46, range 2—3) per multiple removal. On average, 27.4 attacks were selected per removal (± 20.7, 1—100), against 16.7 attacks per control event (± 0.9). A total of 6024 attacks was selected across all removals, against 3387 attacks (± 173) on average per control set.

#### 2.3.4 Hypotheses

We explored three possible non-exclusive alternative hypotheses:

1. **Hypothesis 1**: The lethal removals decreased attack intensities by removing problem individuals (Linnell et al., 1999).
2. **Hypothesis 2**: The lethal removals decreased attack intensities by decreasing local wolf density (Bradley et al., 2015). Relationship between wolf density and attack intensity is also modulated by local richness of wild prey (Petridou et al., 2019).
3. **Hypothesis 3**: The lethal removals decreased attack intensities at small analysis scales and increased attack intensities at intermediate and large analysis scales, by altering the territorial limits of remaining wolves (Brainerd et al., 2008).

We tested the three hypotheses by applying separately the BACI method on different sets of lethal removals defined according to different criteria.

Our first test was made by comparing results from the single removals (n = 188) and from the multiple removals (n = 35). We predicted that a similar decrease of attack intensity observed for both datasets could be in favor of Hypothesis 1, *i.e.* the removal of only one attacking wolf was sufficient to reduce attack intensities. We predicted that a stronger decrease of attack intensity for the dataset of multiple removals could be in favor of Hypothesis 2, *i.e.* attack intensity was correlated to wolf density. Following Hypothesis 3, we predicted that the dataset of multiple removals could more likely support this hypothesis than the dataset of single removals, because risks of territorial alterations increase with the number of killed wolves within a pack (Cassidy et al., 2023).

Our second test was made by dividing the dataset of single removals according to the 3 removal administrative types, as explained in section 2.1 (Figure 1). In practice, the higher the attack intensity in an area, the higher the authorized effort to remove a wolf, from *Simple defense* (n = 65), through *Reinforced defense* (n = 71), and finally *Hunting* (n = 52). Note, however, that the three subsets still concerned removals of only one wolf.

Following Hypothesis 1, we predicted that the *Hunting* subset could present a weaker decrease of attack intensity than the two other subsets, or even no effect, because *Hunting* removals could kill wolves outside of a context of livestock defense, *i.e.* wolves that could not have been responsible for attacks. Following Hypothesis 2, we predicted that the efficiency of removals of *Simple defense*, *Reinforced defense* and *Hunting* to decrease attack intensity would be decreasing following this order, *i.e.* efficiency of removing one wolf decreases with attack intensity.

Our third test was made by dividing the dataset of single removals according to removal location (Figure 4). We selected the 8 zones with at least 10 removals (Figure 4): *PNRPA* (n = 33), *Mercantour* (23), *Vercors* (18), *Trois-Evêchés* (18), *Vanoise* (15), *Préalpes de Digne* (12), *Ecrins* (12) and *Canjuers* (11).

**Figure 4.**
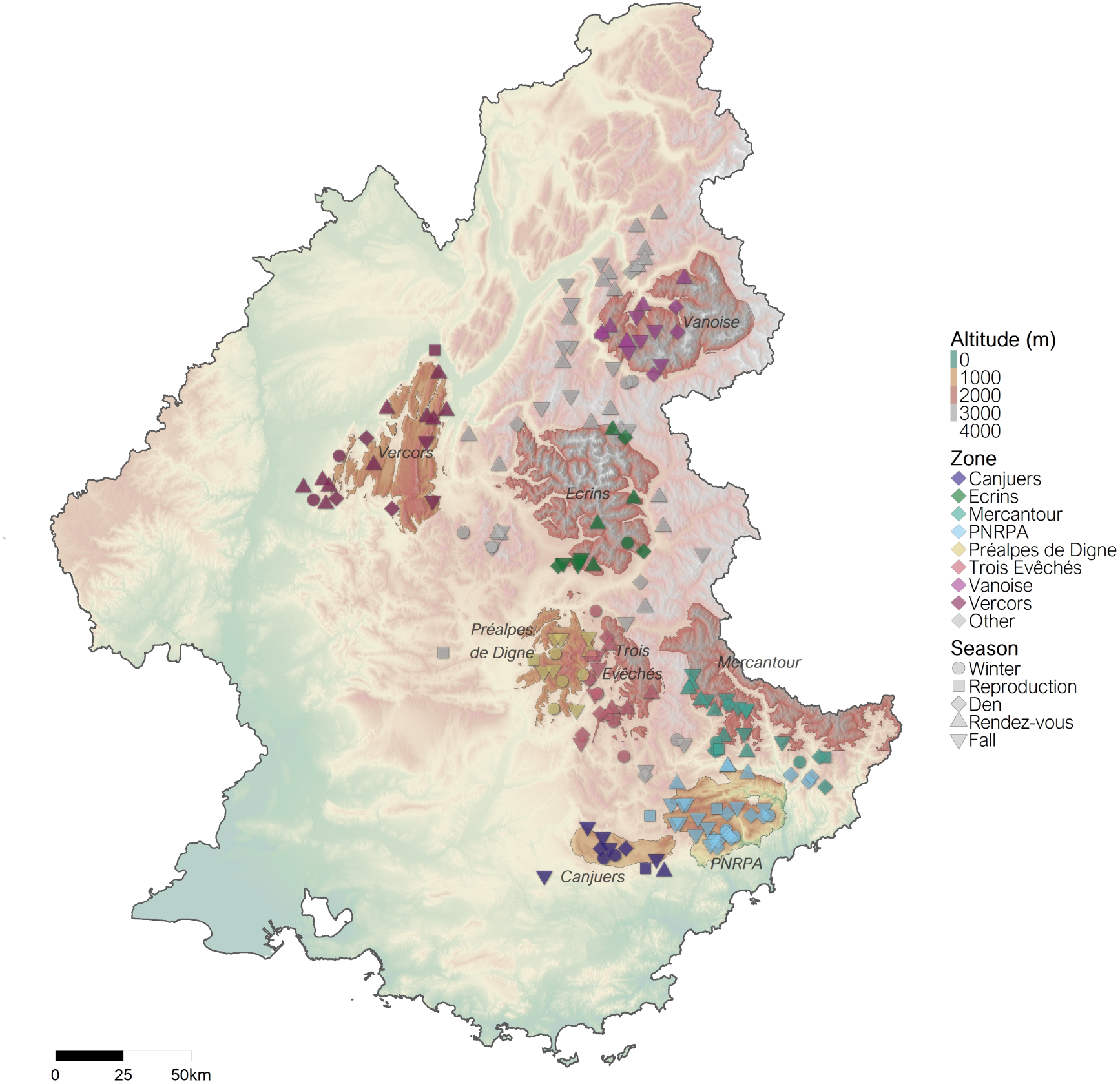
Topography of the study area. Points represent single removals according to their zone and the wolf season during which they occurred. The 8 zones selected for the analyses of the geographic subsets are highlighted.

Following Hypothesis 2, we predicted the weaker decrease of attack intensity for the 3 subsets with the earliest wolf recolonization (*i.e.* starting before 2000), which could result in high local wolf density: *Mercantour*, *Préalpes de Digne* and *Vercors*. Conversely, we predicted the strongest decrease of attack intensity for the 2 subsets with the latest wolf recolonization (*i.e.* after 2015): *Vanoise* and *Ecrins*. Eventually, we predicted intermediate results for the 3 subsets with wolf recolonization starting between 2000 and 2010: *Canjuers*, *Trois Evêchés* and *PNRPA*. Still in line with Hypothesis 2, we also predicted that removals applied in the least mountainous areas, *Canjuers* and *PNRPA*, could be among the least efficient subsets to decrease attack intensity, because wolves could rely more heavily on livestock in these two areas due to a weaker wild ungulate species richness (Petridou et al., 2019). Following Hypothesis 3, we predicted that the subsets with the earliest wolf recolonization would less likely support this hypothesis because their high density of pack territories could decrease the possibilities of territorial changes.

Our fourth test, focused on Hypothesis 2, was made by dividing the dataset of single removals according to the wolf seasons during which removals occurred (Figure 4): *Winter* (November — January, n = 29), *Reproduction* (February — March, 8), *Denning* (April — June, 33), *Rendez-vous* (July — August, 56) and *Fall* (September — October, 62). We predicted that the subsets of *Rendez-vous* and *Fall* could be the least efficient seasonal subsets to decrease attack intensity because of the higher local wolf density due to the presence of growing pups. Similarly, the subsets of *Reproduction* and *Denning* could be the most efficient because of the higher risks of reproduction failure due to removals, leading to reduced pack sizes, but also because of the presence of young wild ungulates during the *Denning* period that were alternative easy prey. We predicted that the subset of *Winter* could present intermediate efficiency, because there are less wolves in winter than in summer, while there are less young wild ungulates, making livestock particularly attractive if available (Mattioli et al., 2011).

Our fifth and final test, focused on Hypothesis 2, was made by dividing the dataset of single removals according to the sex of the killed wolf when known, *Male* (n = 83) and *Female* (n = 78). The removals of the breeding females during spring may greatly impact pup survival, and thus pack size and local wolf density. However, during fall and winter, breeder’s presence also increases pup survival regardless of breeder’s sex (Ausband et al., 2017a). In our dataset, most of female removals were done during summer and fall, and only 7 females over 78 were declared as being reproductive the year they were killed (annual reproductive status of males was unknown). Therefore, we did not predict different results between the two subsets. Breeder removal has greater impacts on the pack stability than other removals (Cassidy et al., 2023), however we could not test Hypothesis 3 because the distribution of removals was too unbalanced between female reproductive status or between age class (177 adults, 24 subadults and 16 pups).

### 2.4 Implementation

The analysis was conducted in R 4.4.2 (R Core Team, 2023). We used the R packages *tidyverse 2.0.0* (Wickham et al., 2019) and *sf 1.0-12* (Pebesma and Bivand, 2023) to handle the datasets. We used the R package *spatstat 3.2-1* (Baddeley et al., 2015) to simulate point patterns. We adapted the function *kde2d.weighted* from *ggtern 3.4.2* (Hamilton and Ferry, 2018) to separate the kernel estimates before and after the removal date. We used the package *osmdata 0.2.5* (Padgham et al., 2017) to extract limits of PNRPA and of Canjuers military zone. We used the packages *geodata 0.5-8* (Hijmans et al., 2023), *ggfx 1.0.1* (Pedersen, 2022a), *ggnewscale 0.5.2* (Campitelli, 2023), *ggpubr 0.6.0* (Kassambara, 2023), *metR 0.14.1* (Campitelli, 2021), *patchwork 1.3.1* (Pedersen, 2022b), *RColorBrewer 1.1-3* (Neuwirth, 2022), *scales 1.3.0* (Wickham and Seidel, 2022), *tidyterra 0.7.2* (Hernangómez, 2023) and *tidytext 0.4.3* (Silge and Robinson, 2016) for data visualization. The datasets (with anonymized coordinates), codes and results are available online (Grente et al., 2025).

## 3. Results

### 3.1 General spatio-temporal distribution of attacks around lethal removals

Spatio-temporal distribution of attacks around lethal removals took various forms according to the datasets or subsets (Figures 5, S1). However, we could extract two features common. First, removals were generally preceded by high attack intensities at their locations, meaning they were applied in response to attacks, in accordance with French legislation. Second, attacks tended to mostly aggregate below 2 km from removal locations, regardless of time. Therefore, the attack process was generally a persistent phenomenon at these locations, either before or after removals, while generally presenting several interruptions in time.

**Figure 5.**
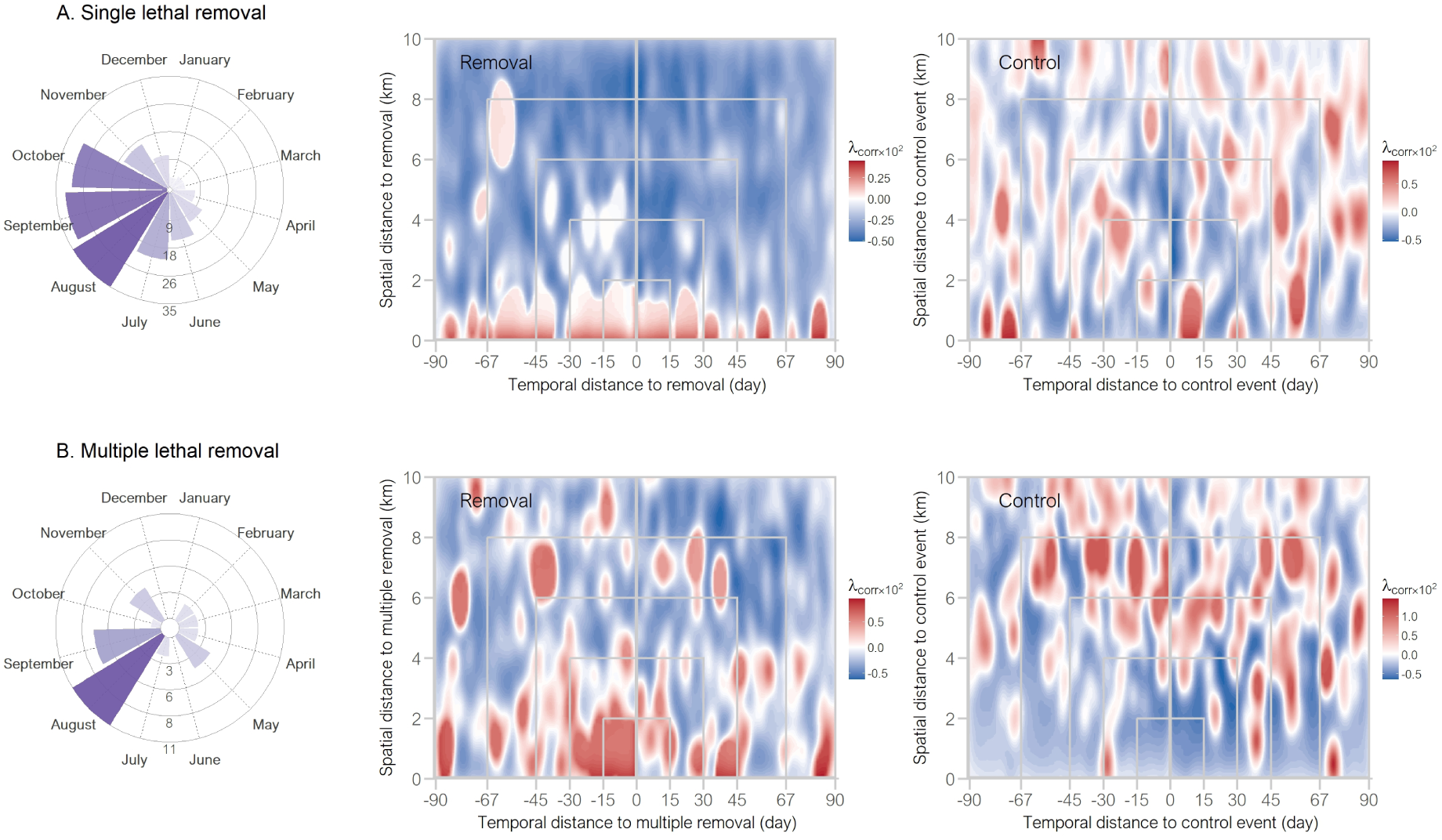
Corrected attack intensities around lethal removals (middle) and around the first set of control events (right), for the dataset of single removals (A) and of multiple removals (B). Each panel includes the monthly distribution of the dataset. Grey rectangles delimit the five nested scales used to calculate trends in Figures 6, 8 and 9.

### 3.2 General effect of removals

The trends calculated for the dataset of single removals (*i.e.* one killed wolf) and for the dataset of multiple removals (*i.e.* on average killed 2.29 wolves) were significantly negative across the five nested analysis scales (Figure 6).

**Figure 6.**
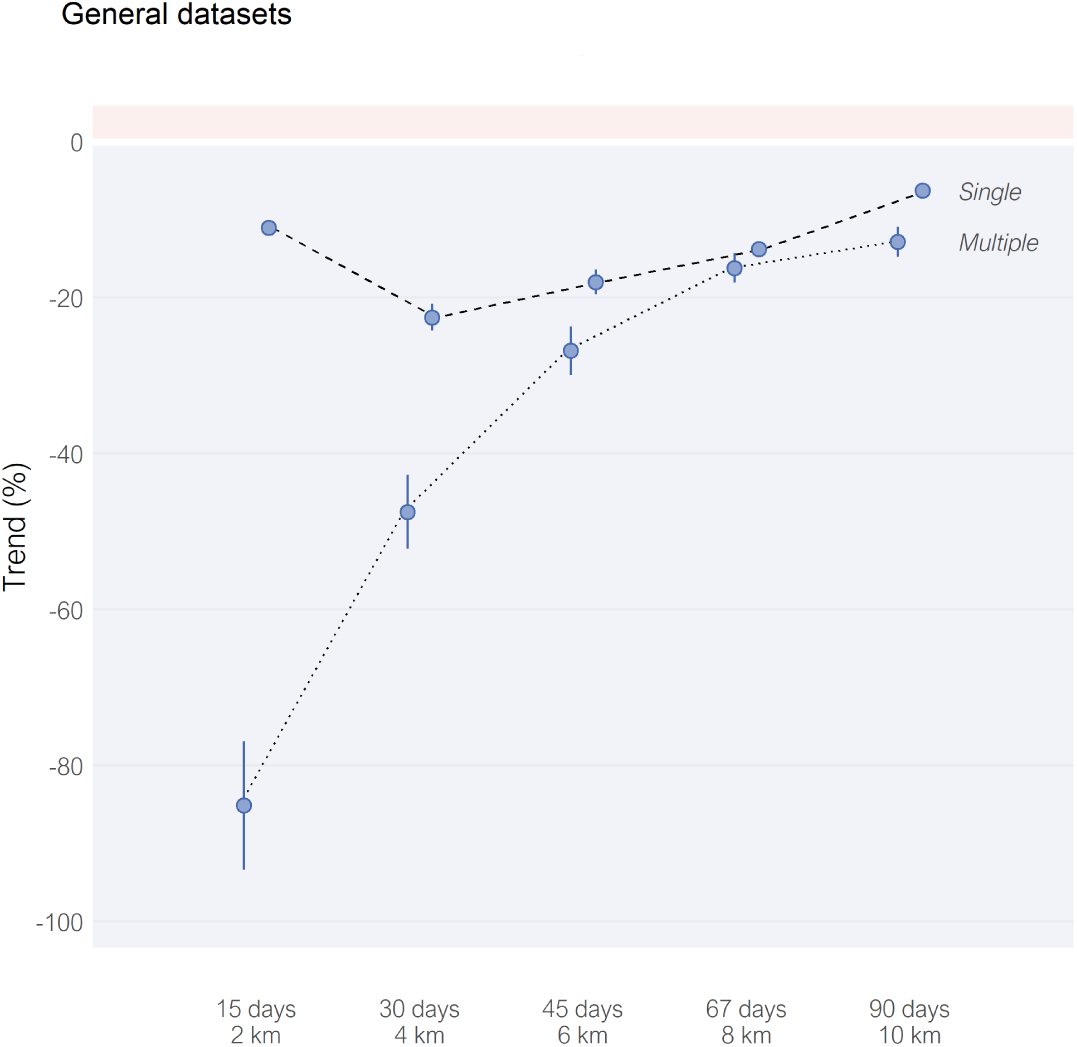
Trends of attack intensities after the applications of single removals and multiple removals, at five nested scales. Points are mean estimates, alongside the confidence intervals (range 83.4%–89.8%). All trends are significant. Trends were corrected for livestock presence.

The trend of attack intensity after single lethal removals ranged from -22.5% (−24.2% – -20.8%) to - 6.3% (−7.15% – -5.38%) according to the analysis scale (Figure 6), suggesting the lethal removals of one wolf moderately decreased the attack intensities.

By comparison, the trend of attack intensity after multiple removals ranged from -85.1% (−93.4 – - 76.9%) to -12.8% (−14.7 – -10.9%) according to the analysis scale (Figure 6), suggesting the lethal removals of 2 or 3 wolves very strongly to moderately decreased the attack intensities.

### 3.3 Effect of the analysis scale

The analysis scale had a limited effect on the trend for the dataset of single removals (Figure 6). At fixed spatial scale, the decrease of attack intensity remained moderate over 90 days after the removal (Figure 7A). However, the short periods after single removals, below 15 days, could show non-significant trends.

**Figure 7.**
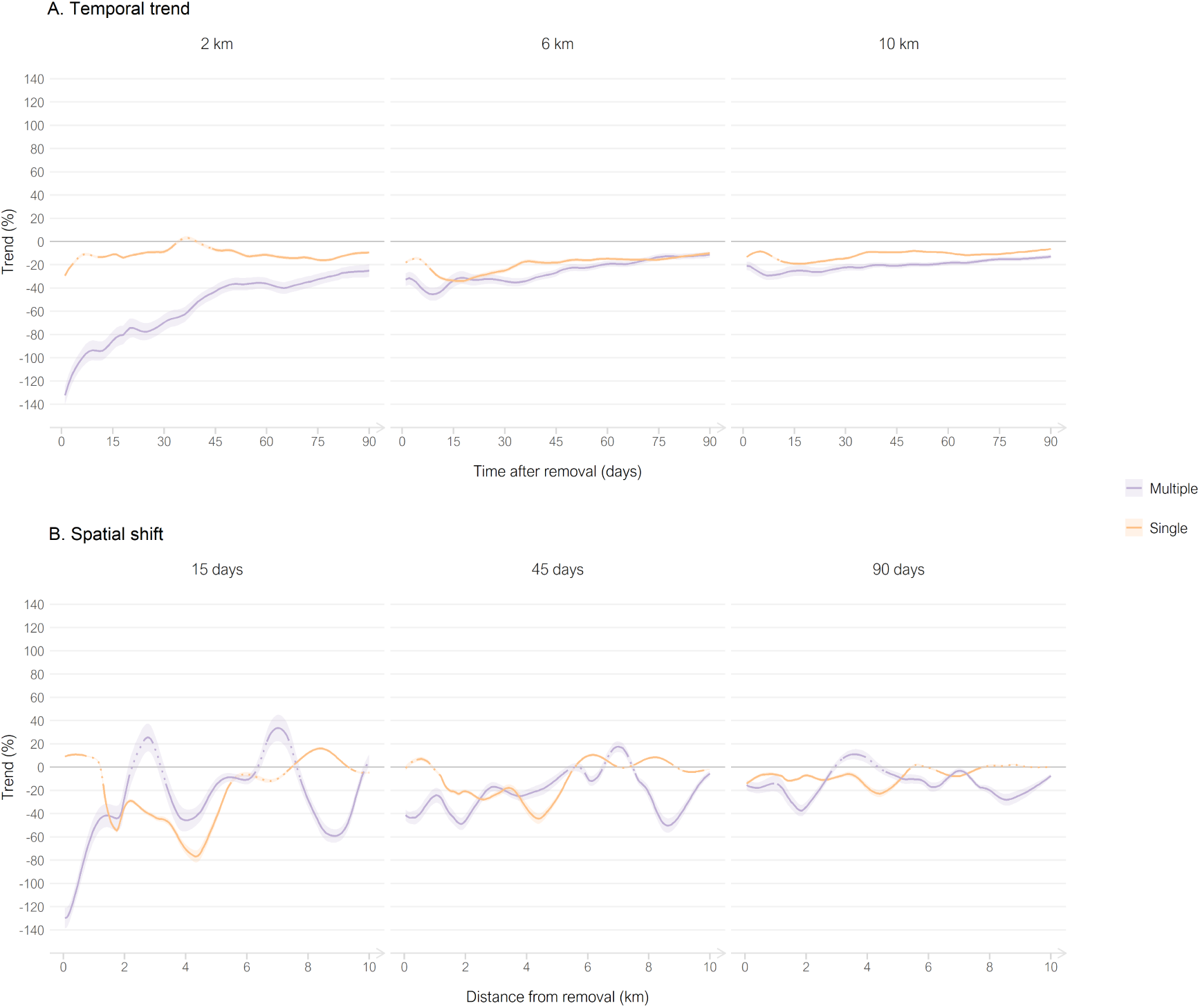
Trends of attack intensities after the applications of single removals (orange) and of multiple removals (purple), alongside continuous temporal (A) and spatial (B) scales, for three fixed distances and periods, respectively. Lines are mean estimates, ribbons are the confidence intervals (range 83.4–90.4%). Dotted lines are not significant trends. Trends were corrected for livestock presence. Note that temporal trends are calculated as nested periods (e.g. from 0 to x days, over 0–2 km for first panel), while spatial shifts are trends calculated for specific distances (e.g. at the xth km, over 15 days for first panel).

On the contrary, the analysis scale had an important effect on the trend for the dataset of multiple removals (Figure 6). Over 0–2 km, the trend distinctly increased with time (Figure 7A), going from the strongest decrease at day 1 (−124% – -140%) to the smallest decrease at day 90 (−30.4% – - 19.5%). The very strong effect of reduction of multiple removals disappeared and became moderate when focusing on larger spatial scales, joining the trend of single removals over the entire time scale (Figure 7A). This suggests that the very strong reduction induced by multiple removals was mostly localized to the targeted pasture and to its closest neighbors over a period of more than one month.

Spatial shifts of attacks after removals were not obvious (Figure 7B). For most distances, trends were negative, regardless of the fixed temporal scale. Nevertheless, some significant positive trends could appear at certain distances (e.g. around the 3^rd^ and 7^th^ km, 15 days after multiple removals) but they had not the magnitude of negative peaks. This might suggest that a minor part of the attack intensities shifted towards other pastures after removals.

### 3.4 Effect of the correction for livestock presence

The correction for livestock presence had a limited effect on the trends for the dataset of single removals, but reinforced the magnitude of negative trends for the dataset of multiple removals (Figures 6, S2, 7, S3). In other words, raw attack intensities decreased after multiple removals, but this decrease took on even more significance in light of the fact that most pastures were still in use after these multiple removals, and could therefore have been attacked but were mainly not.

This difference between single and multiple removals could be explained by the fact that single removals were more evenly distributed between summer and fall, while multiple removals were mainly applied in August (Figure 5). The dataset of single removals (n = 188) was also larger than the dataset of multiple removals (n = 35). Therefore, the needs for a correction would be less important for a large dataset with various underlying livestock distributions.

### 3.5 Effect of subsets

The main effect of moderate decrease of single removals concealed disparities between single removals. When dividing this large dataset according to removal’s location, removal’s season, removal’s administrative type or the sex of the killed wolf, we could observe very different trend profiles across the subsets (Figures 8, 9).

**Figure 8.**
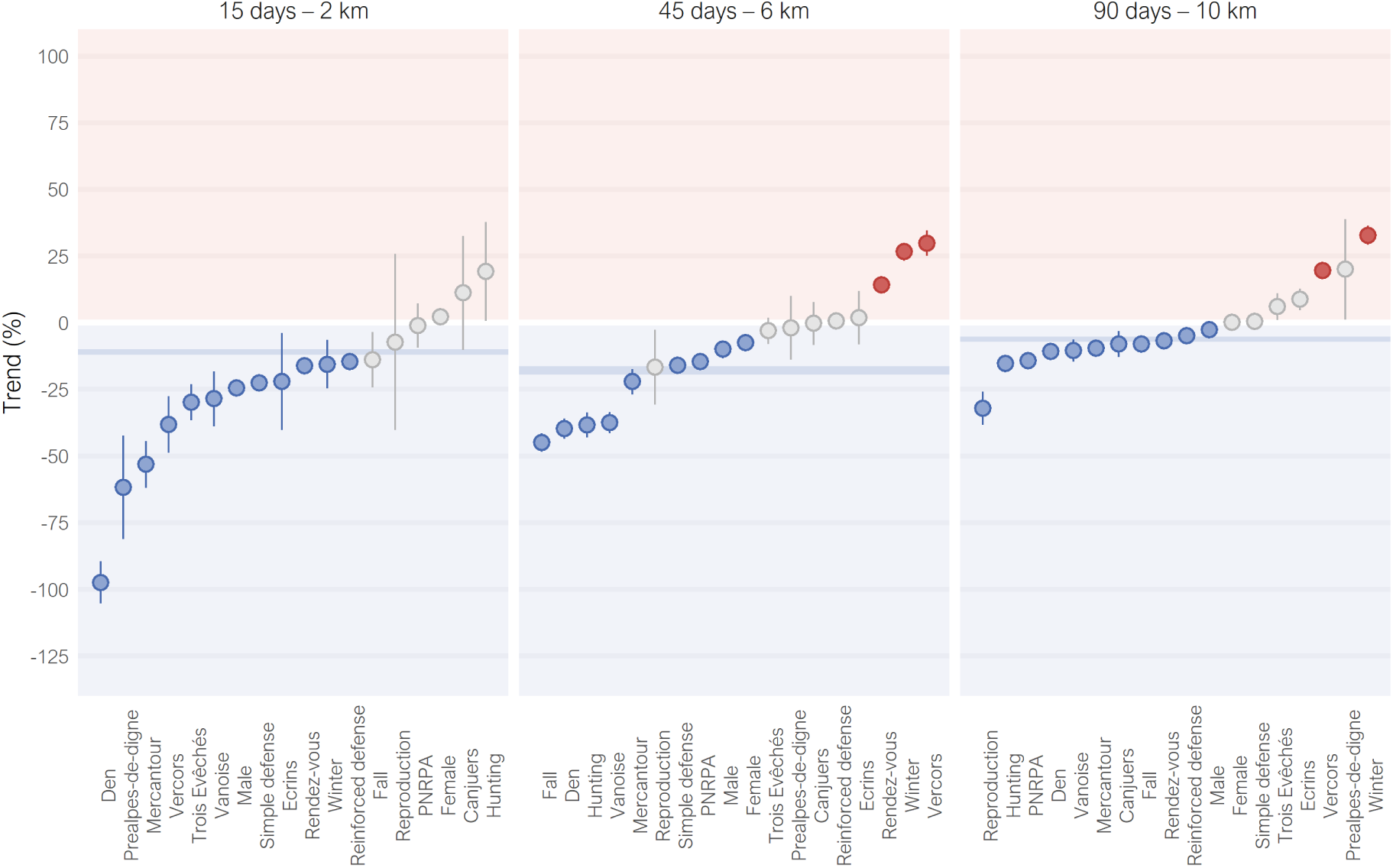
Distributions of the trends of attack intensities for the 18 subsets of single removals at three nested scales. Points are mean estimates, alongside the confidence intervals (range 83.4%–89.8%). Trends in grey are not significant. Trends were corrected for livestock presence. Confidence intervals of the entire dataset of single removals are in background in dark blue.

**Figure 9.**
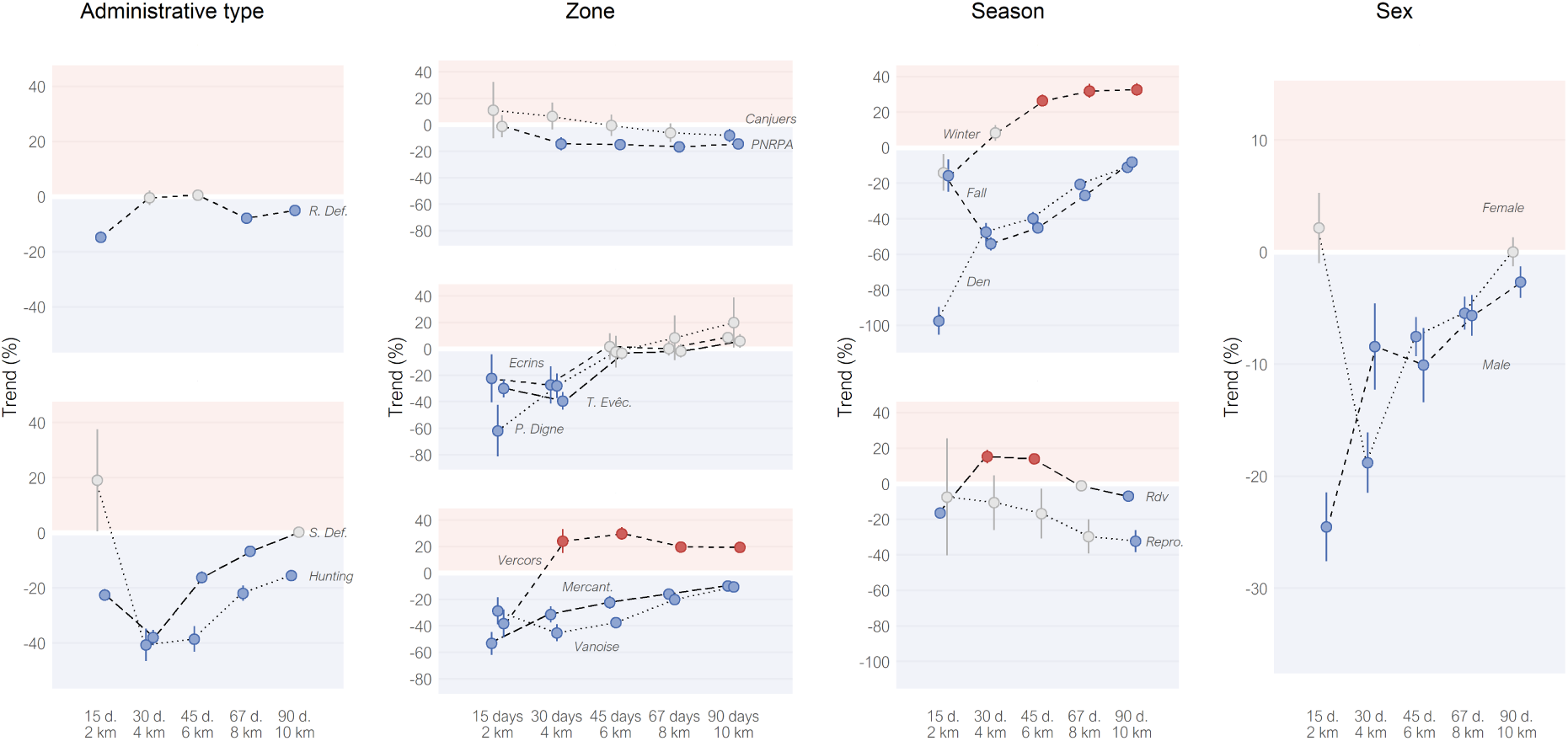
Trends of attack intensities after the applications of single removals, divided into 18 subsets, at five nested scales. Points are mean estimates, alongside the confidence intervals (range 88.5%–89.8%). Trends in grey are not significant. Trends were corrected for livestock presence. ‘R. Def’ and ‘S. Def’ for Reinforced and Simple defense subsets; ‘T. Evêc.’, ‘P. Digne’ and ‘Mercant.’ for Trois Evêchés, Préalpes de Digne and Mercantour subsets; ‘Rdv’ and ‘Repro.’ for Rendez-vous and Reproduction subsets.

At the smallest nested scale (*i.e.* over 15 days and 0–2 km), we observed a large gradient of significant negative trends (Figure 8). Besides 6 subsets that presented moderate decreases as the general dataset of single removals (from -5% to -24%), 3 subsets presented very strong decreases (below -50%) and 3 others strong decreases (from -49% to -25%). When increasing the analysis scale, most subsets showed similar results to the general dataset of single removals (Figures 8, 9).

In addition, there were three subsets that showed significant positive trends (*Vercors*, *Winter* and *Rendez-vous*), while their trends at the first nested scale were significantly negative (Figure 8). Besides the increase of attack intensity, spatial shifts could be suspected for these subsets (Figure S4B). The correction for livestock presence for these subsets played an important role to reveal most of these positive trends, that could be not significant or even negative without correction (Figures S5, S6). In these cases, this meant that the raw decrease of attack intensities was in fact expected given the fact that most pastures were not in use anymore after the removals, and that the attack intensity actually increased when taking this bias into account. This was especially true for the Winter subset, because its removals occurred at the limit between an intense and calm grazing period (Figure S1).

These deviations from the general dataset of single removals, whether through strong decreases or increases after single removals, revealed particularities that could happen in some situations, but did not represent the main effect that we could expect after single removals. Except the *Rendez-vous* subset, the sizes of the subsets with these deviations were indeed below the average subset size (*n*=37.3).

### 3.6 Exploring the variability of the effect

This variability among subsets could be explored through the results to the tests of hypotheses we described in section 2.3.4.

Hypothesis 1 (“*lethal removals decreased attack intensities by removing problem individuals*”) was not supported. Single removals did not show similar trends as multiple removals, and removals from the Hunting subset were not the least efficient among its category (Figures 6, 9).

Hypothesis 2 (“*lethal removals decreased attack intensities by decreasing local wolf density*”) received partial support (Figures 6, 9). As expected, multiple removals presented a better efficiency to decrease attack intensity than single removals. However, the prediction that removal efficiency decreased with the intensities of attacks preceding removals, tested with the subsets of the administrative types, was not verified by the results. Results were mixed regarding the geographic and season subsets. The subsets of *Canjuers* and *PNRPA* were the only subsets that did not show strong negative trends after removals, in line with the possible higher reliance of wolves on livestock there, due to a weaker wild ungulate species richness. However, the order of efficiency of geographic subsets expected according to the historical wolf recolonization did not clearly support the hypothesis. The subsets of *Denning* and *Rendez-vous* verified the predictions of Hypothesis 2, but not the subsets of *Reproduction* and *Fall*. Contrary to the initial prediction, the subset of *Winter* did not present intermediate efficiency, but instead showed positive trends at the largest scale. This could match with part of the prediction, where we suggested that wolves may heavily depend on livestock at this season. Eventually, the absence of large trend differences between the subsets of female and male removals was consistent with Hypothesis 2.

The results for Hypothesis 3 (“*lethal removals altered wolf territorial limits*”) also received partial support. As predicted, the dataset of multiple removals more supported this hypothesis than the dataset of single removals (Figure 7B). However, two of the three subsets with the earliest wolf recolonization (*Vercors* and *Préalpes de Digne*) presented signs of spatial shift (Figure S4B), contrary to the prediction.

## 4. Discussion

We evaluated the effects of lethal control of wolves on attack intensity distribution through kernel density estimation. Our method provided an efficient visualization of attack intensity distributions in the form of heatmaps over continuous spatial and temporal scales, corrected for sheep presence. The resulting attack intensity trends showed the differences in the attack intensities before and after lethal removals (intra-site comparison) and with comparable areas without removals (BACI design).

The aim of our study was to generalize the spatio-temporal patterns of attack intensity surrounding lethal removal applications, without concealing the potential diversity of these patterns across the French Alps that could result from different situations. This is why we compared trends of single lethal removals and trends of multiple lethal removals, but we also divided single lethal removals into 18 subsets according to the application conditions (geographical, seasonal, administrative) or to the removal results (sex).

### 4.1 Assessment of lethal removal efficiency

The distribution of the attacks was usually structured by space, with strong spatial aggregations of attacks, especially close to the locations of the lethal removals. This suggested that some locations may have been more subject to attacks than others, including at a local scale (< 10 km).

Removals of several wolves (2.29 wolves on average) were more efficient to reduce attacks than lethal removals of one wolf, especially at low analysis scales. Multiple removals reduced the attack intensity by 85% over 15 days and 2 km, against 11% for single removals. Results of both datasets were getting closer with increasing analysis scales. If we focus on the pastures targeted by multiple removals and their closest neighbors (< 2 km), the reduction of attack intensity remained very strong or strong during 89 days. These results are consistent with the results from Bradley et al. (2015), where removal efficiency increased with the number of killed wolves in the northern Rocky Mountains of the USA.

The effect of analysis scale was also visible for the 18 subsets of single lethal removals. The moderate decrease of single lethal removals transformed into a large gradient of trends at the subset level. Generally, the smaller the area and the time period surrounding the lethal removals, the stronger the decrease of attacks after removals. Three subsets also showed positive trends while their trends at the lowest analysis scale were negative, and showed signs of spatial shifts of attacks.

Our results highlight the importance of accounting for scale in such assessments. Therefore, the observed disparity between the results of studies evaluating wolf lethal control could partly result from the disparity in the analysis scales. If the effect of removals generally weakens with analysis scale, like in our results, the likelier it is to find non-significant results when assessing removal effects over large spatio-temporal scales. For example, Kutal et al. (2023) found no significant effect of wolf removals on attacks in Slovakia, but analyzed the datasets over 711 km² (314 km² here) and over one year (3 months here) without intermediate scales.

Differences in analysis scales cannot be the only explanation for differences between studies, as Santiago-Avila et al., (2018) found non-significant effects while using a progressive spatial scale comparable to ours. Besides analysis scales, other methodological choices, linked to the analyzed datasets, may explain these differences. Here, our dataset was very unbalanced in favor of attacks, with a ratio of 1 removal for 27 attacks, against 1 for 7 in Santiago-Avila et al., 2018. Thus, the risk of recurrence of a single attack was the chosen metric in several studies with datasets from the USA (Bradley et al., 2015; Harper et al., 2008; Santiago-Avila et al., 2018), whereas we chose to focus on the number of attacks (*i.e.* intensities).

### 4.2 Limits of the analysis

We expect these results to reasonably reflect the effects of lethal removals on wolves. Thus, the lethal removals should have generally decreased attack intensities by temporarily decreasing local wolf density or by altering the distribution or the predation behavior of the remaining wolves.

However, as in every study, confounding factors may interfere. Here, the main risk was to confound the effects of lethal removals on wolves with the effects of livestock distribution dynamics in space and time. We corrected our analysis for livestock distribution based on a pastoral census in the Alps that gave the precise limits of pastures and their months or seasons in use (Dobremez et al., 2016). This type of regional census requires a substantial amount of work. Fine-scale livestock data as provided by this census is therefore generally missing from analyses on predation, including for other parts of France (*e.g.* Gastineau et al., 2019). When integrated, livestock data is generally at large, administrative scales (*e.g.* DeCesare et al., 2018; Merz et al., 2025; Šuba et al., 2023). Thus, only two comparative studies (Allen, 2015; Conner et al., 1998) and two correlative studies (Blejwas et al., 2002; Wagner and Conover, 1999) that evaluated lethal control effects on canids corrected their analysis for livestock presence at the pasture level.

However, despite the quality of the livestock census we used, this may not have been enough to prevent the main confounding risk for our study. Annual changes in the use of pastures since the census may have occurred. In addition, temporary changes in livestock presence are common during a grazing season, as livestock owners and shepherds modulate the use of pastures by their flocks to optimize grazing according to environmental conditions. Among these conditions, pressure due to wolf predation or attempts of predation may encourage livestock owners to remove flocks from some pastures or to reinforce preventive measures. Data on precise husbandry practices, such as preventive measures, were out of the scope of this census. Hence, resulting potential variations in livestock distribution and protection, even small and temporary, may drive changes in attack distribution that would not have been considered in the census and therefore in the livestock correction of our analysis. To a lesser extent, variations in the spatio-temporal distributions of wild ungulates may also impact the distribution of attacks on livestock.

In spite of the limits of the livestock dataset, the intra-site comparison design should have helped to reduce the confounding risks, especially at the smallest scale. Because wolves move over large territories, we acknowledge that focusing on changes happening only in very close proximity to removals could not be enough to evaluate removal effects (Treves et al., 2016). To improve the statistical power of our analysis, especially at the larger analysis scales, we doubled the intra-site comparison with an inter-site comparison, therefore applying a BACI design, evaluated as robust by Christie et al. (2019). The main limit of this study design is to use appropriate control sites. In our case, this was a challenging task because lethal removals in France were generally authorized and applied in attack hotspots, creating unbalanced initial attack intensities between sites with and without removals. Nevertheless, the buffers of control events had on average 16.7 attacks (± 0.9) against 26.3 (± 20.7) for the removal buffers, suggesting that control sites were also subject to attacks.

### 4.3 Underlying mechanisms

If the confounding risks were as low as possible, we can conclude that lethal removals generally decreased attack intensities, with only a few situations leading to other results (non-significant or increase). We explored several hypotheses to understand the underlying mechanisms leading to these results. Our results did not support the hypothesis of problem individuals (Linnell et al., 1999). In addition, for most analyzed subsets, attacks persisted after removals at the location of the removals, including at high intensity, even though most removals targeted wolves approaching livestock or attempting attacks. This result suggested that attacks on livestock were more a structural phenomenon than the outcome of some problem individuals.

The better efficiency of multiple removals compared to single removals clearly supported the hypothesis of a reduction of attacks through the temporary decrease of local wolf density. The initial local wolf density might also explain the gradient of attack responses between the subsets of single removals. However, the support to this hypothesis was less clear-cut, probably because our characterization of the subsets was too rough and simplistic to describe local contexts surrounding lethal removals.

Therefore, the gradient of attack responses after removals observed between subsets of single lethal removals remains largely unexplained. Besides effects on wolf density, removals may have altered the distribution or the predation behavior of the remaining wolves, as spatial shifts could be suspected for some subsets, like in Santiago-Avila et al. (2018). However, the persistence of attacks at the removal’s locations in most situations, even at lower intensities, challenges the hypothesis of pack dissolution (Elbroch and Treves, 2023). Indeed, settlement of new wolves at the same location was unlikely to be as fast, even in the dense population of the Western Alps (Marucco et al., 2023).

### 4.4 Implications for management and research

Concluding that the analyzed removals were an efficient management tool requires to define efficiency itself. As in most studies on the subject, we considered that the significant decreases of attack intensities after removals met expectations of efficiency, at least for the analyzed scales.

However, other definitions are possible, depending on the magnitude of the reduction, the spatio-temporal scales at which the effects apply, and whether or not removals threaten wolf population viability (*e.g.* Lennox et al., 2018). Expectations regarding removal efficiency also depend on the benefits and costs of wolf’s conservation, that can be very different across people (Linnell et al., 2010). Thus, research results about management actions can be differently interpreted by people (Hodgson et al., 2019). This is why clear management objectives of lethal control must be defined through open dialogue, in view of ecological responses like here, but also by considering socioeconomic dimensions and wolf population viability (Lennox et al., 2018; Linnell et al., 2010).

We showed that the analyzed lethal removals of wolves generally helped to reduce attack intensities within the targeted pastures and their neighbors. By contrast, some particular situations did not lead to this result (*e.g.* removals within *Vercors*, *Canjuers,* or during July-August). Considering our results are still valid in today’s situation, this variability may be a thorn in manager’s side, as wolf removals will probably lead to attack reduction, but not with certainty. Given the strong impact lethal control can have on wolf population (Grente et al., 2024), the next step for research would be to move forward from patterns to mechanisms. This understanding is necessary to provide the keys for managers for the most efficient application of lethal control in terms of attack reduction, and therefore for a better public acceptance of this tool.

We could only briefly explore the mechanisms of the effects of lethal removals on wolves and therefore on their attacks. An advanced analysis would have required to have access to pack compositions and space use by wolves at fine spatio-temporal scale, which were unavailable. Collection of these datasets on wolves are particularly difficult on large areas, considering the secrecy of wolves. A trade-off would be to deploy the substantial means required to collect such datasets in a small study area (*e.g.* one wolf territory and its neighboring packs) for a few years, ideally replicated at different locations. Effects of lethal control on wolves and their attacks may then be evaluated through promising methods such as animal social network analysis, which was applied to study the reaction of other social animals to human-induced removals (*e.g.* Downing et al., 2023; Goldenberg et al., 2016). Working at small scales would also facilitate a more precise collection of livestock presence and husbandry practices.

Variability is inherent to any ecological process (*e.g.* human avoidance by wolves, Ferreiro-Arias et al., (2024); wolf mortality, Milleret et al., (2025)). Predation behavior is no exception. A few fine-scale datasets on wolves and livestock would not necessarily explain all the observed removal effects on attacks. A solution would be to upgrade the analysis of observed data by simulations (Railsback et al., 2025). In this sense, we simulated removal effects on wolf population dynamics and on attacks at the scale of a mountain range in the Alps (Grente et al., 2024), but our theoretical model would greatly benefit from a combination with social network analysis based on observed data at small scale.

## 5. Conclusion

The lethal management strategy implemented in France from 2011 to 2020—primarily involving the removal of single wolves in defense situations with protected livestock—can be expected to have produced a moderate reduction in attack intensity following removals. This reduction was greatly amplified when removing multiple wolves. Our results partly supported the hypothesis of removal effects through reduction of local wolf density but also through changes in the distribution of the remaining wolves. However, contextual factors (e.g., geographical or seasonal variations) could lead to deviations from this general pattern. Future research should focus on understanding wolves’ responses to removals—such as changes in pack composition, space use, and predation behavior— to better interpret the observed variability in attack trends post-removal.

## Supplementary material

Three supplementary materials are available online, on Zenodo (https://doi.org/10.5281/zenodo.17231870).

File 1. Supplementary information (S1–S3)

File 2. Supplementary figures (S1–S9)

File 3. Supplementary tables (S1–S2)

## Acknowledgements

We are indebted to the public officials in charge of recording attack claims and lethal removals. We are grateful to the regional administration of Auvergne-Rhône-Alpes for the access to the databases of claims and of wolf lethal removals. We are also grateful to INRAE (previously IRSTEA) for the access to the pastoral census. We greatly thank Marion Valeix, John Linnell and Michel Meuret for their valuable comments regarding the first draft of this study as part of the PhD dissertation of Oksana Grente. We greatly thank Aaron Bott, Vincenzo Gervasi, Peter Sunde and an anonymous reviewer for their feedbacks, that greatly helped us to improve the manuscript. We also thank members of the scientific council of the National Plan of Action for wolves and livestock breeding activities for their feedback and advices. The analyses were performed on the Core Cluster of the Institut Français de Bioinformatique (IFB). A preprint version of this article has been peer-reviewed and recommended by PCI Ecology (https://doi.org/10.24072/pci.ecology.100752). We greatly thank Tomas Willebrand for overseeing the peer review process.

## Data, scripts and code

Data, scripts and code are available online, on Zenodo (https://doi.org/10.5281/zenodo.17231870) and on gitlab (https://gitlab.com/oksanagrente/Kernel_wolf_culling_attacks_p).

## Conflict of interest disclosure

The authors declare that they comply with the PCI rule of having no financial conflicts of interest in relation to the content of the article.

Olivier Gimenez and Simon Chamaillé-Jammes are recommenders of the managing board of PCI Ecology.

