## Supplementary information for "Spatio-temporal dynamics of attacks around deaths of wolves: A statistical assessment of lethal control efficiency in France"

---

*Supplementary Information S1: Attributing attacks to pastoral units*

The attack dataset showed 59% of consistency with the dataset of pastoral units, meaning that 59% attacks were within pastures that were in use according to the pastoral census. The other attacks could be outside pastoral units, or could have occurred in months for which pastoral units were not supposed to be in use according to the pastoral census. These mismatches could result from changes in the spatial or temporal use of pastoral units, from errors during the attack record (*e.g.* geolocation approximation), or because attacks occurred in non-indexed pastoral units.

When attacks were within pastures and when difference between the month of their dates and the sets of grazed months of their pastures did not exceed three months (9% of total attack dataset), we only updated the pastoral census by adding the months of the attacks to the corresponding pastures. Otherwise, we relocated attacks to their closest pastures within a 500 meters radius by favoring pastures with the smallest difference between their sets of grazed months and the months of the attacks. When these differences exceeded 3 months, or if no pastoral unit could be found within a 500 meters radius around the attacks, we excluded these attacks (13% of total attack dataset). Relocations were done at the closest point of the pastoral units to the original attack locations (19% of total attack dataset). When necessary, we added the months of the relocated attacks to the sets of grazed months of their new pastoral units. The final dataset of attacks had 19 302 events.

Contrary to attacks, lethal removals did not require to be located within pastoral units, as hunting did not necessarily occur within pastoral units.

---

*Supplementary Information S2: Spatio-temporal distribution of attacks around removals*

This section explains in greater details how we selected the attacks for each removal through buffers (*1. Construction of buffers*), how we applied a spatial correction to each attack (*2. Reducing the number of spatial dimensions*), and how we simulated attacks according to pastoral distribution (*3. Correction for livestock presence*).

1. *Construction of buffers*

We attributed to each lethal removal a spatio-temporal 'observed buffer', which can be visualized as a cylinder centered on the lethal removal event (Figure S8.A). The circle base of the cylinder corresponded to the spatial dimensions (X, Y) and had the role to scan for attacks within the pastoral space around the lethal removal. The height of the cylinder corresponded to the temporal dimension, the left and right parts scanning for attacks that had occurred before or after the lethal removal, respectively.

The observed buffer was composed solely of the pastoral units where attacks could occur. Therefore, the buffer was sparse, and remained empty where there was no pastoral unit around the removal event according to the pastoral census.

We set the radius of the cylinder at 10 km, in order to encompass more than one wolf territory radius, estimated at 7–8 km in France through telemetry and genetic tracking (Duchamp et al. 2012). Therefore, ripple effects, if any, could be observed up to 10 km from the lethal removal. The pastoral units overlapping the buffer were truncated to the buffer.

We set the height of the cylinder to 180 days, meaning that all attacks occurring 90 days before and 90 days after the day of the lethal removal were part of the buffer. Thus, we analyzed the effect of lethal removal on a full season if the removal event occurred in the middle of the season, or on two successive seasons if not. The day of the removal was not part of the buffer, because we could not know if an attack recorded at the day of the removal occurred after of before the removal, having no information about the hour of both attacks and removals.

Attacks were not homogeneously distributed across space and time (Grente et al. 2022). Increasing the radius or the height of buffers compressed the attack distribution towards the centers of buffers (*i.e.* at small spatial and temporal distances), and prevented an optimal interpretation of the results.

We ensured that the buffers did not overlap geographic areas or periods without data about attacks. We removed 84 lethal removals occurring at less than 10 km from the terrestrial limits of the study area or occurring after the 1$^{st}$ October 2020, eventually leaving 278 analyzed lethal removals.

Most buffers overlapped in space and in time, with a total of 86% of buffers overlapping with at least another one. However, only a part of removal events were simultaneously close in space and time (Figure S9), with 82 removals separated by less than 5 km and less than 25 days. In these cases, we considered that the risks of interaction between removals were high. We pooled these removals into 36 groups, with each group corresponding to a set of removals (hereafter, `a multiple removal`). For the special case where two removals had no closely overlapping buffers, but both closely overlapped with a third removal, we considered these three removals as part of the same group of removals. On average, 2.28 wolves (± 0.45, range 2—3) were killed per multiple removal.

We attributed a single observed buffer to each multiple removal, of 10 km radius, of 180 days height and centered on the centroid of the removals of the group. To compare the attack intensity before and after each multiple removal, we excluded the attacks recorded between the date of the first removal of the group and the date of the last removal of the group. The exclusion period lasted 7.5 days on average (± 8.38, range 0—27). In cases where removals occurred the same day (28% of multiple removals), only the date of the removals was not part of the buffer, as previously.

We ended up with 232 observed buffers, including 196 buffers of single removals and 36 buffers of multiple removals, corresponding to a total of 6110 attacks. On average, an observed buffer included 26.3 attacks (± 20.7, range 0—100).

2. *Reducing the number of spatial dimensions*

Methods for the analysis of interactions of several datasets with three dimensions are yet to be developed and may suffer from the curse of dimensionality (Wikle et al. 2019). Therefore, we followed standard geostatistical practices and reduced the two spatial dimensions of the observed buffers to one, by computing the Euclidian distance of each attack to the center of the buffer. In other words, we converted the spatial circle of buffers to a line. The temporal dimension corresponded to the difference in days between each attack and the date of the removal of the buffer. In the case of a multiple removal, we used the first (or last) date of the group if the attack occurred before (or after) the exclusion period. Thus, each attack had two coordinates $(i, j)$ corresponding to the spatial $i$ and temporal $j$ distances from the center of the observed 2D-buffer (Figure S8.B).

Nevertheless, converting a 2D-circle into a 1D-line always induces a spatial bias that gives more representation of points farther away, simply because the circle perimeter increases linearly with circle radius. Therefore, we should have applied a spatial correction of $1/i$ to each attack of an observed buffer, with $i$ the Euclidian distance of the attack to the buffer center. However, the buffers were delimited by pastoral units that were not homogeneously distributed around the buffer center. In other words, the spatial bias did not increase linearly with the radius, but according to the perimeter fraction that intersected with pastoral units. For example, the spatial bias could be low at the furthest distances of the buffers if pastoral units were almost non-existent at these distances.

To incorporate pastoral units to the spatial correction, we sliced up each spatial buffer into 200 rings $r$ of width 50 m. We calculated $pa_r$, the total area of pastoral units of each ring $r$. We then attributed to each attack from the observed 2D-buffer the inverse of the pastoral area of its ring, $1/pa_r$ (Figure S8.C). Because attacks could only be located in pastoral units, $pa_r$ of rings containing attacks could not be null.

3. *Correction for livestock presence*

For each removal, we corrected for livestock presence by randomly simulating attacks with a distribution defined according to the locations of pastoral units of the buffer and their grazed months. The simulated dataset corresponded to the expected distribution of attacks if they only depended on livestock presence.

For each month $m$ represented in the observed 3D-buffer, we calculated the pastoral area $pa_{r,m}$ that was used per ring $r$ during month $m$ (Figure S8.D). Because the information about pastoral use was monthly and not daily, we duplicated the unique column of $pa_{r,m}$ into the number of days of this month that were part of the buffer (Figure S8.E).

This resulted in a matrix of 200 x 180 dimensions (one row per ring, one column per day). The matrix coordinates were similar to the coordinates *(i, j)* of the observed attacks, with a slightly coarser spatial resolution of 50 meters (*i.e.* ring width) and the same temporal resolution of 1 day.

We converted the matrix into a two-dimensional pixel image. We generated a random point pattern, with the pixel image acting as the probability density of the points. We generated the same number of points as the number of observed attacks of the buffer. We extracted the coordinates *(i, j)* of each point (hereafter, 'simulated attack'), corresponding to its spatial and temporal distances to the buffer center (Figure S8.F). We also attributed the spatial correction $1/pa_r$ for each simulated attack (Figure S8.G). We repeated the simulation 1000 times. Thus, for each observed buffer, we ended up with 1000 2D-buffers of simulated attacks (hereafter, 'simulated buffers').

---

*Supplementary Information S3: Kernel density estimation*

1. *Attack intensity estimation*

For each set of removals (defined in section 2.3.4 of the main text), all the observed 2D-buffers of the removals of the set were aggregated into a unique 'observed set buffer' (Figure 2). The point of coordinates $i = 0$ and $j = 0$ of the set buffer then represented the aggregated spatial location of all the removals of the set, and the day before the removals (*i.e.* the day of the removals was not part of the buffer).

We estimated the observed attack intensities $\lambda_{obs}$ of the set buffer by applying an anisotropic Gaussian kernel density estimation (Figure 2). The method consisted in dividing the observed set buffer into pixels, and in estimating the observed attack intensity of each pixel of coordinate ($i$, $j$). We used the same pixel resolution as previously, *i.e.* 50 meters and 1 day. A Gaussian function (*i.e.* 'kernel') was centered over each pixel, and could spread over adjacent pixels. The spread of the kernel was determined by the bandwidth parameters. We used two separate bandwidth parameters for spatial distance and temporal lag, of 500 meters and 2 days. The choice of the bandwidth parameters was *ad-hoc*, and met the needs of adopting a resolution close to the pixels while smoothing attack intensities enough in order to obtain interpretable results. The attack intensity of a pixel $\lambda_{obs}(i,j)$ was then calculated by summing the value of the attacks within its kernel. The value of an attack was equal to its spatial correction $1/pa_r$, in order to correct for the reduction in the spatial dimensions. Thus, the higher the spatial corrections of the attacks within a kernel, the higher the attack intensity estimate of the related pixel, $\lambda_{obs}(i,j)$.

We also considered the spatio-temporal distributions of attacks before removals ($j \in [-90, 0]$ and after removals ($j \in [1, 90]$) as two separate distributions by appropriately right- and left-truncating kernels in the time dimension for pixels occurring before and after the date of removals, respectively. Thus, kernels that were centered before removals could not spread over spatial coordinates located after removals, and vice versa.

We corrected kernel estimates for outliers by capping the estimated observed attack intensities $\lambda_{obs}(i, j)$ to $Q_3 + 1.5\times(Q_3 - Q_1)$, where $Q_1$ and $Q_3$ were the first and third quantiles of the whole estimated distribution of $\lambda_{obs}$ of the set buffer (Walfish 2006).

Similarly, we aggregated the first simulated 2D-buffer for each removal of the set into a unique 'simulated set buffer'. We repeated the aggregation for the 999 other sets of simulated buffers. We estimated the simulated attack intensities $\lambda_{sim}(i,j)$ for each of the 1000 simulated set buffers. We then calculated the mean simulated attack intensities $\bar{\lambda}_{sim}(i,j)$ which corresponded to the expected attack intensities given pastoral use.

To correct the observed kernel estimation for livestock presence, we calculated the corrected intensities $\lambda_{corr}(i, j)$ as:

$$\lambda_{corr}(i,j) = \lambda_{obs}(i,j) - \bar{\lambda}_{sim}(i,j) \quad (1)$$

Thus, a value of $\lambda_{corr}(i, j)$ lower, equal or higher than zero indicated that the observed attack intensity at coordinates *(i, j)* was lower, equal or higher, respectively, than the attack intensity expected according to pastoral use.

2. *Trends of attack intensity after removals*

We calculated the trends of observed or corrected attack intensities at different spatio-temporal scales *s* as:

$$Y_s = \frac{\sum_{i=1,j=1}^{i=I_s,j=J_s} \lambda'(i,j) - \sum_{i=1,j=-(J_s-1)}^{i=I_s,j=0} \lambda'(i,j)}{I_s \times J_s} \times 100 \quad (2)$$

where $I_s$ and $J_s$ were the number of rows (spatial distance to removals) and columns (number of days before or after removals) of scale *s*, $\lambda'(i, j)$ was the observed or corrected intensity at the coordinates *(i, j)* rescaled over [0,1] for observed intensity or over [-1,1] for corrected intensity by using the minimum and maximum values of $\lambda_{obs}(i, j)$ or $\lambda_{corr}(i, j)$ over the temporal range $-(J_s-1), J_s$. The rescaling allowed trend comparison between sets of removals. The division by $I_s \times J_s$ allowed trend comparison between scales *s*.

We chose 5 nested spatio-temporal scales *s*, where spatial and temporal ranges both increased:

- Scale 1: Trends over 0–2 km and over ± 15 days ($I_1 = 40$; $J_1 = 15$)
- Scale 2: Trends over 0–4 km and over ± 30 days ($I_2 = 80$; $J_2 = 30$)
- Scale 3: Trends over 0–6 km and over ± 45 days ($I_3 = 120$; $J_3 = 45$)
- Scale 4: Trends over 0–8 km and over ± 67 days ($I_4 = 160$; $J_4 = 67$)
- Scale 5: Trends over 0–10 km and over ± 90 days ($I_5 = 200$; $J_5 = 90$)

We also calculated the nested trends by keeping the spatial range fixed while increasing the temporal range by one day, starting at ± 1 day, eventually giving 90 temporal trends. We tested three fixed spatial ranges, $I_1$, $I_3$ and $I_5$. For example, the temporal scale n°28 was calculated over ± 28 days, with $I_1$, $I_3$ or $I_5$.

Eventually, we investigated the potential spatial shift of attack intensities by keeping the temporal range fixed while applying Equation 2 at each spatial coordinate *i* of the spatial range 0-10 km ($i \in [1,200]$), eventually giving 200 spatial trends. We tested three fixed temporal ranges, $J_1$, $J_3$ and $J_5$. For example, the spatial trend n°80 was calculated at the 4$^{th}$ km (and not over 0–4 km), with $J_1$, $J_3$ or $J_5$. We only adjusted the spatial trends by $J_s$ as each spatial trend was calculated over one spatial unit only (*i.e.* 50 meters). Namely, we transformed Equation 2 into Equation 3: $Y_s(i) = (\sum_{j=1}^{j=J_s} \lambda'(i,j) - \sum_{j=-(J_s-1)}^{j=0} \lambda'(i,j)) / J_s \times 100$.

*3. Jackknife samples*

We estimated the standard deviation of each attack trend following removals by using the jackknife method (Maesono 2011). We chose to apply the jackknife instead of the bootstrap method to reduce computational time.

To do so, we split the set of removals into as many jackknife samples as removals. Each jackknife sample consisted of the set of removals minus one of the removals, with each time a different withdrawn removal. We computed the trend Y of the attack intensities $\lambda_{corr}$ for each jackknife sample, following the procedure previously described.

Then, we calculated the standard error of the jackknife samples as:

$$SE_{jack}(\widehat{Y_s}) = \sqrt{\frac{n - 1}{n} \sum_{i = 1}^{n} (\hat{Y}_{-i} - \bar{Y})^2}$$

where *n* is the total number of removals of the set, $\hat{Y}_{-i}$ is the trend of the $i^{th}$ jackknife sample and $\bar{Y}$ is the average trend of all jackknife samples. The standard error of the jackknife samples corresponded to the standard deviation of the trend of the set of removals.

---

*References*

Duchamp, C., Boyer, J., Briaudet, P.-E., Léonard, Y., Moris, P., Bataille, A., Dahier, T., Delacour, G., Millisher, G., Miquel, C., Poillot, C. and Marboutin, E. 2012. A dual frame survey to assess time– and space–related changes of the colonizing wolf population in France. - Hystrix Ital. J. Mammal. 23: 12.

Grente, O., Saubusse, T., Gimenez, O., Marboutin, E. and Duchamp, C. 2022. Wolf depredation hotspots in France: Clustering analyses adjusting for livestock availability. - Biol. Conserv. 267: 109495.

Maesono, Y. 2011. Jackknife. - In: Lovric, M. (ed), International Encyclopedia of Statistical Science. Springer Berlin Heidelberg, pp. 697–699.

Walfish, S. 2006. A review of statistical outlier methods. - Pharm. Technol. 30: 82–88.

Wikle, C., K., Zammit-Mangion, A. and Cressie, N. 2019. Spatio-Temporal Statistics with R. - Chapman and Hall/CRC.
