## Supplementary figures for "Spatio-temporal dynamics of attacks around deaths of wolves: A statistical assessment of lethal control efficiency in France"

### Figure S1

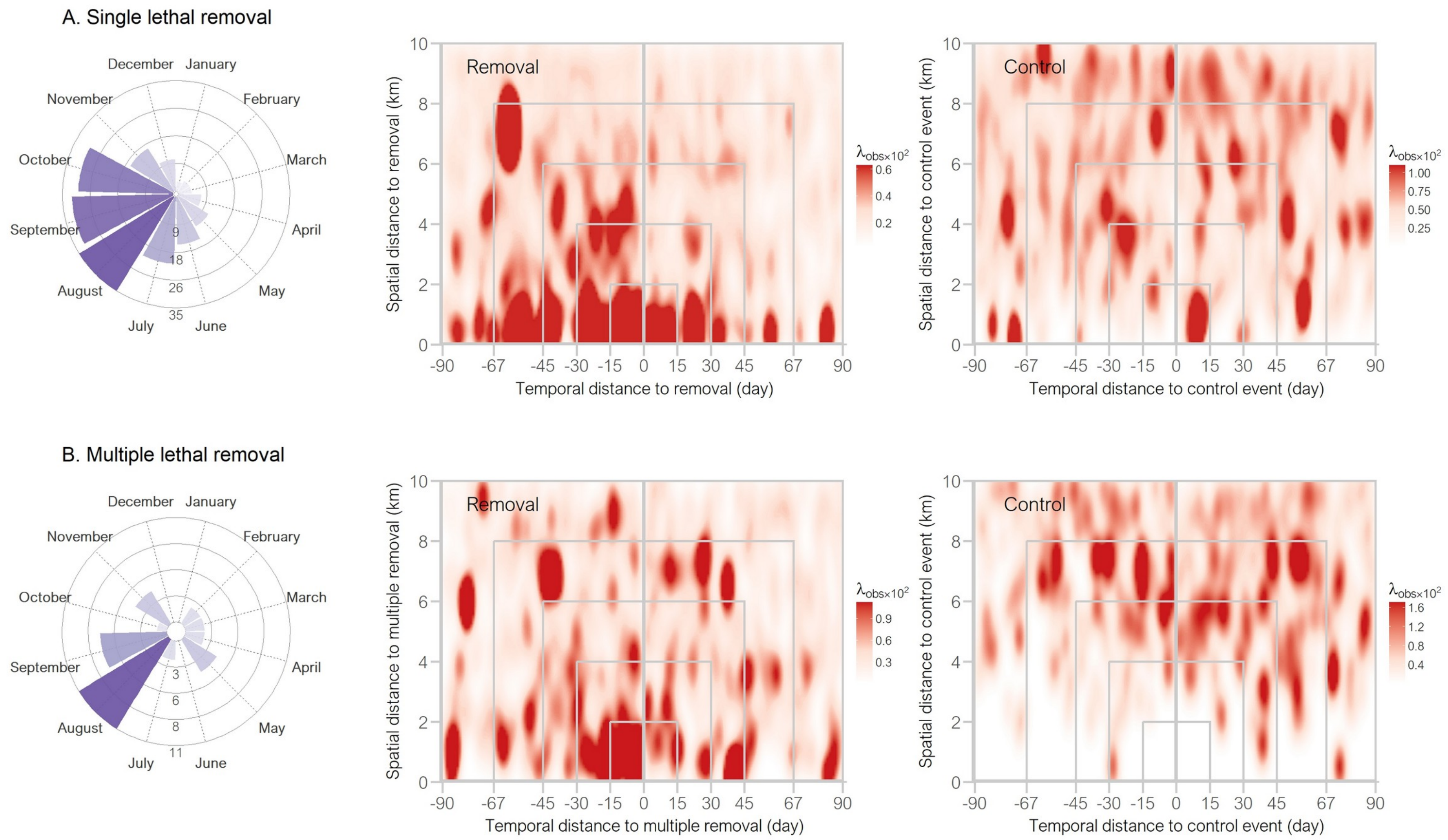


*Figure S1.*

*A-B: Observed attack intensities around lethal removals (middle) and around the first set of control events (right), for the datasets of single removals (A) and multiple removals (B). Each panel includes the monthly distribution of the dataset.*

*C-T: Observed (left) and corrected (right) intensities around lethal removals and around the first set of control events, for the 18 subsets of single removals. Each panel includes the monthly distribution of the dataset.*

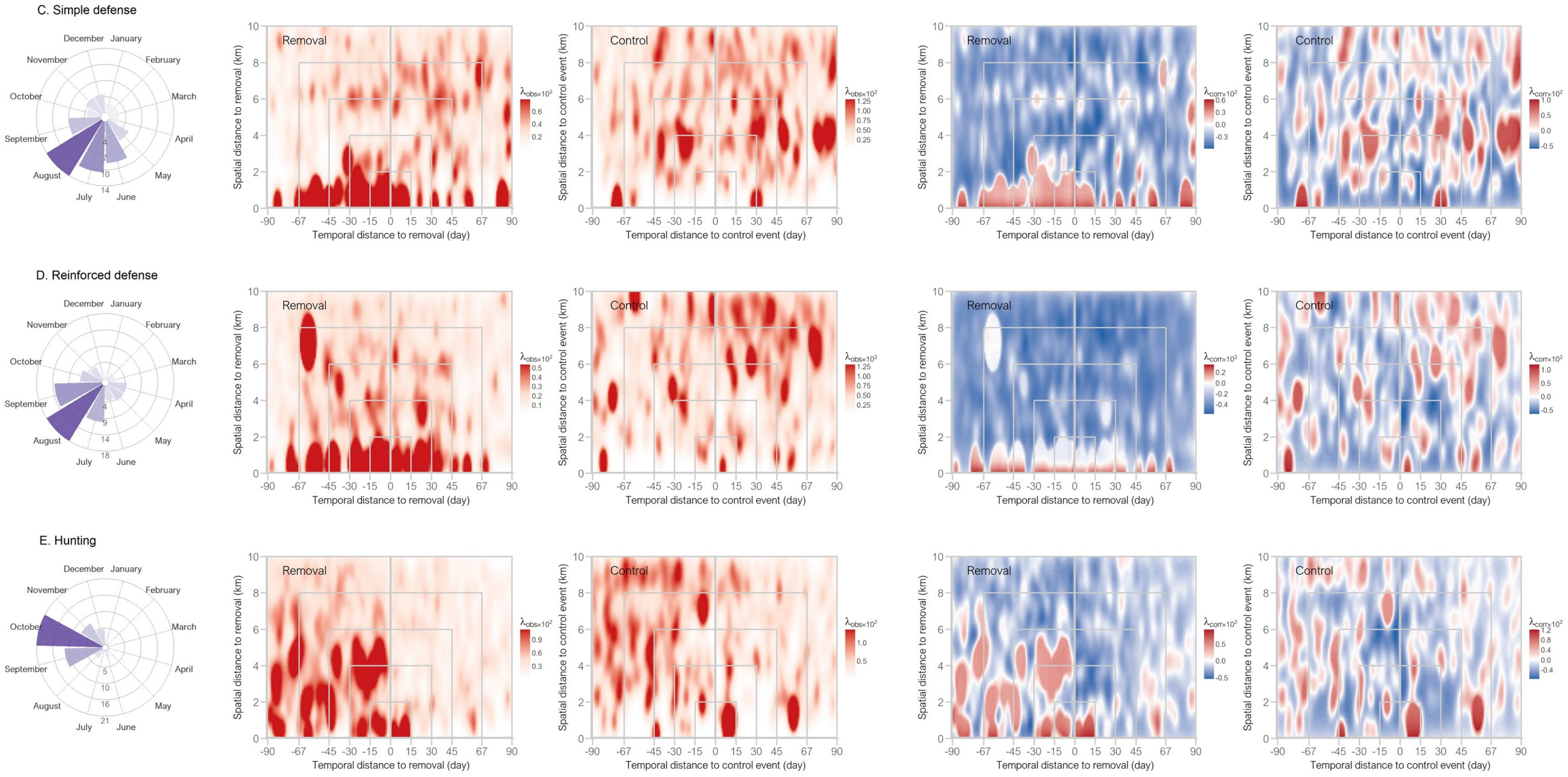

C. Simple defense
D. Reinforced defense
E. Hunting
December
January
February
March
April
May
June
July
August
September
October
November
Removal
Control
Spatial distance to removal (km)
Spatial distance to control event (km)
Temporal distance to removal (day)
Temporal distance to control event (day)
-90 -67 -45 -30 -15 0 15 30 45 67 90
$\lambda_{obs \times 10^2}$
$\lambda_{corr \times 10^2}$

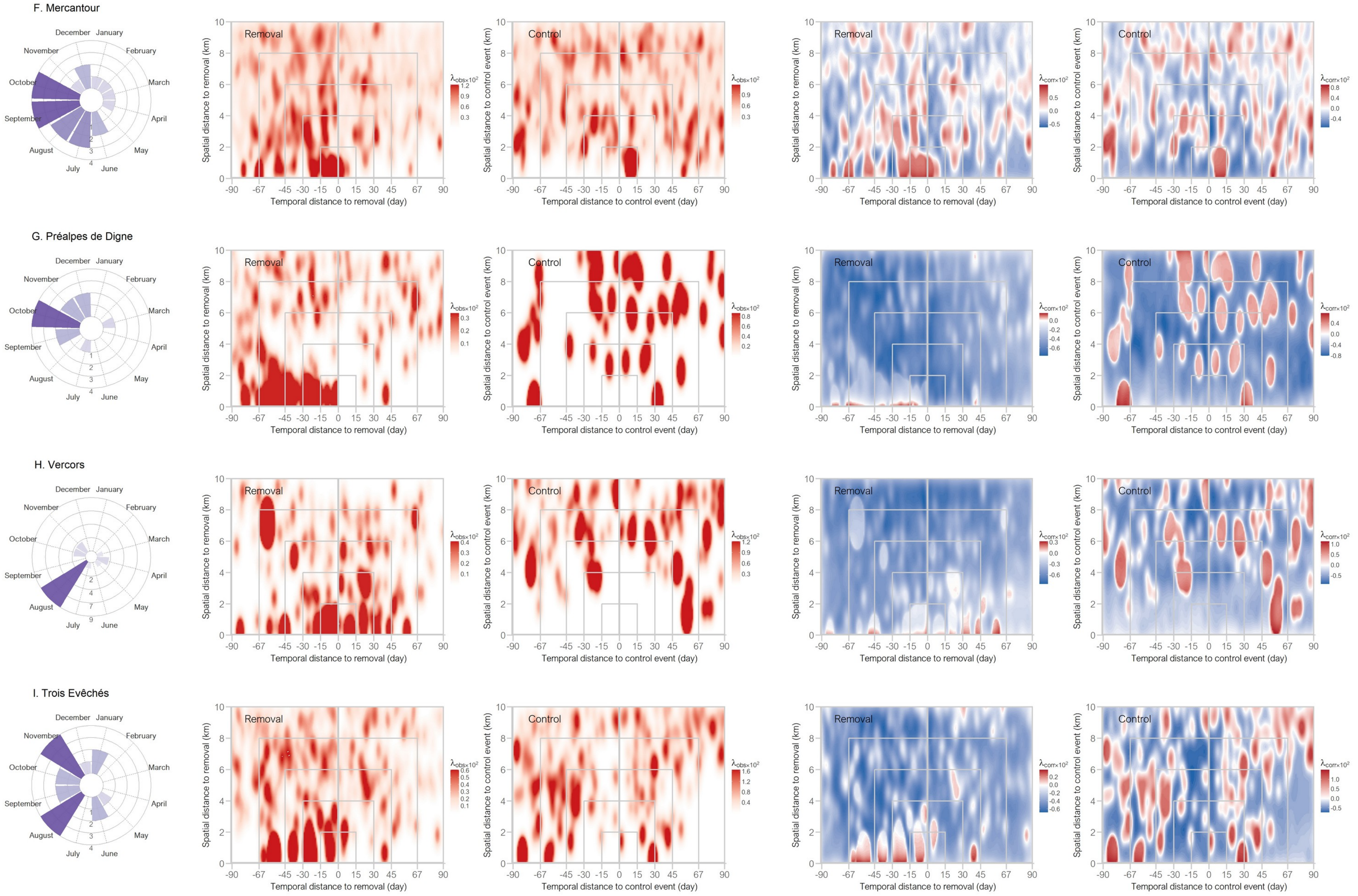
F. Mercantour
G. Préalpes de Digne
H. Vercors
I. Trois Evêchés
December
January
February
March
April
May
June
July
August
September
October
November
Removal
Control
Spatial distance to removal (km)
Spatial distance to control event (km)
Temporal distance to removal (day)
Temporal distance to control event (day)
-90
-67
-45
-30
-15
0
15
30
45
67
90
λobs×10²
λcorr×10²

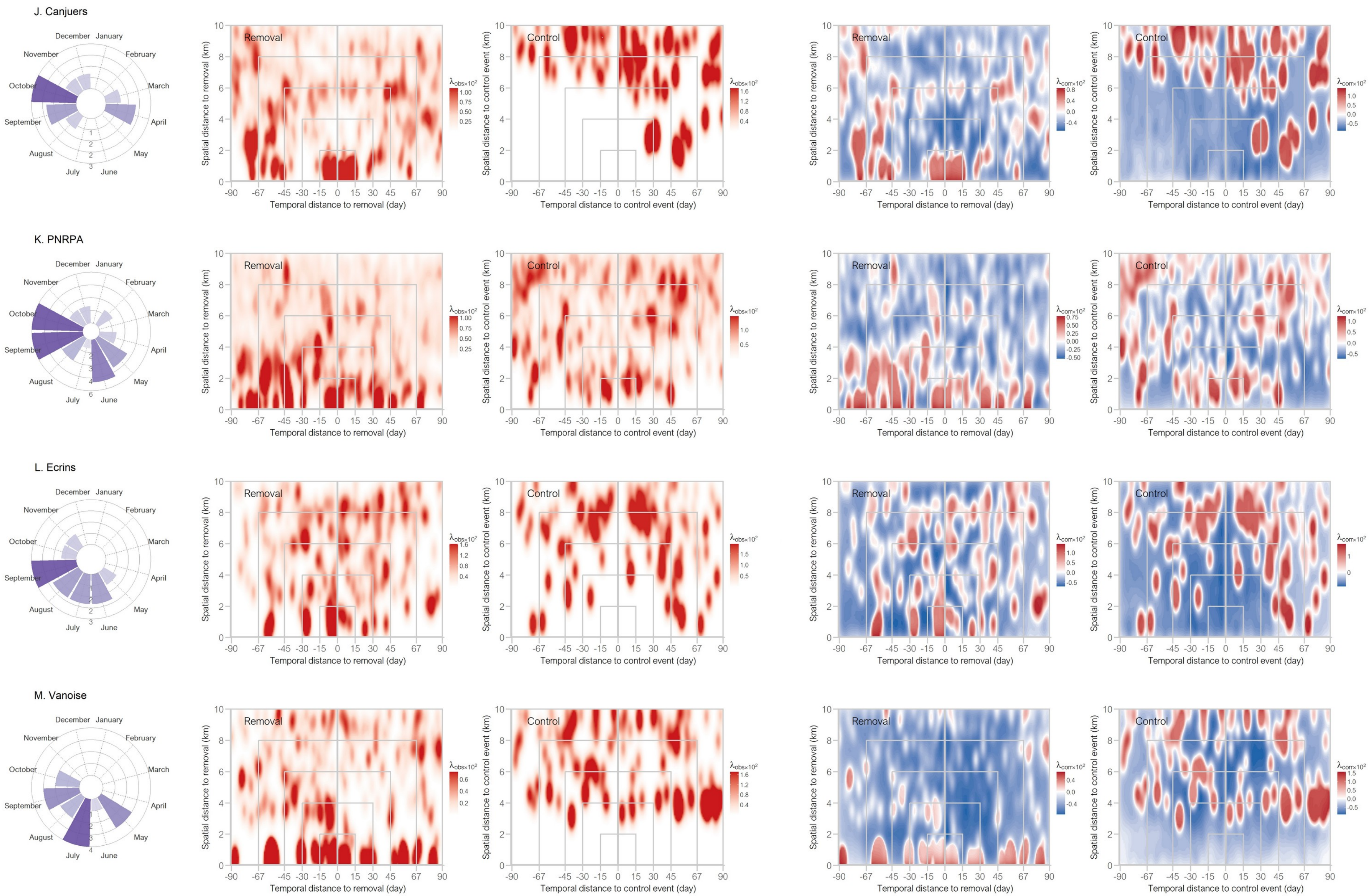
J. Canjuers
K. PNRPA
L. Ecrins
M. Vanoise
December
January
February
March
April
May
June
July
August
September
October
November
Removal
Control
Spatial distance to removal (km)
Spatial distance to control event (km)
Temporal distance to removal (day)
Temporal distance to control event (day)
-90 -67 -45 -30 -15 0 15 30 45 67 90
$\lambda_{obs}\times10^2$
$\lambda_{corr}\times10^2$

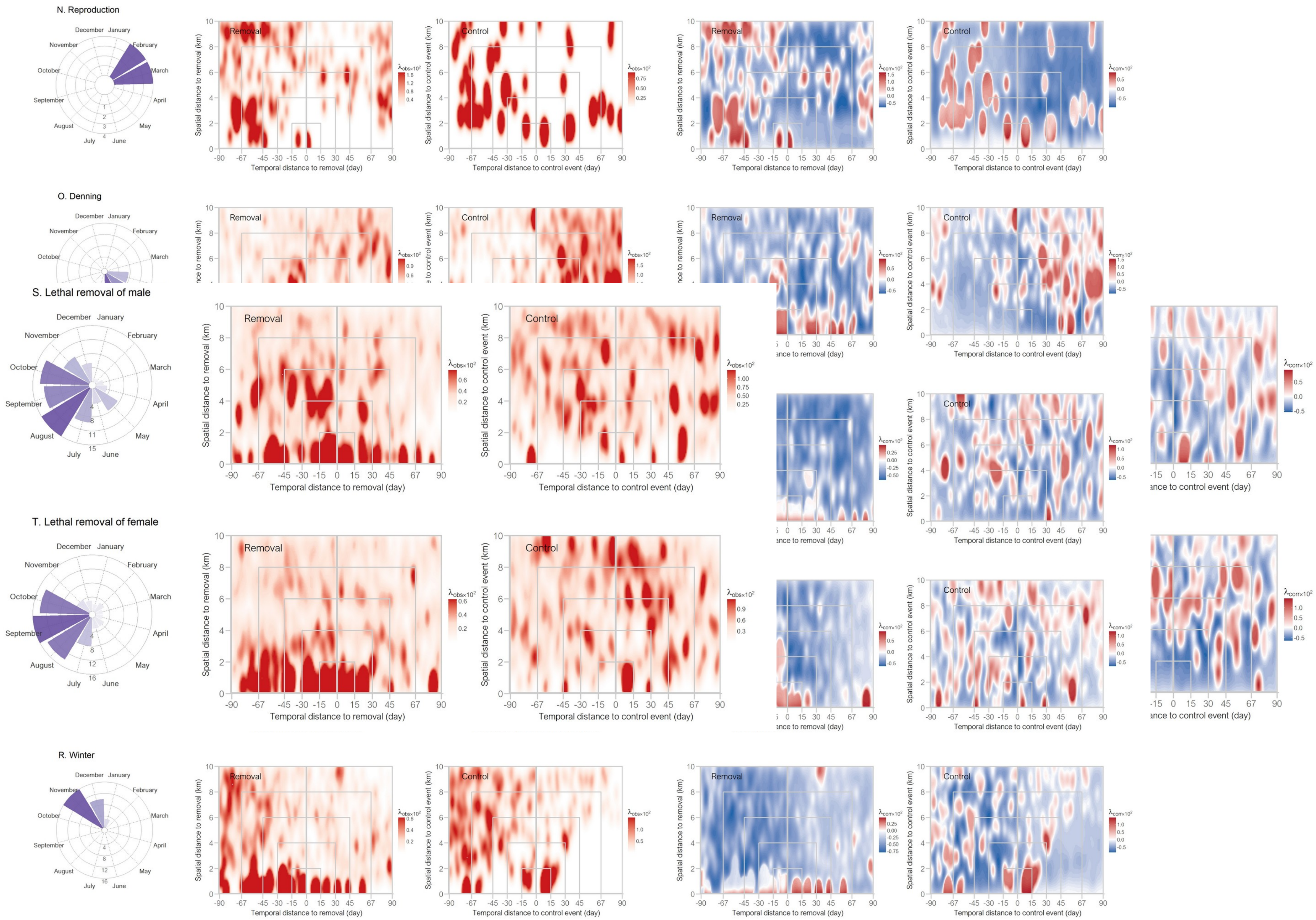
N. Reproduction
O. Denning
S. Lethal removal of male
T. Lethal removal of female
R. Winter
December
January
February
March
April
May
June
July
August
September
October
November
Removal
Control
Spatial distance to removal (km)
Spatial distance to control event (km)
Temporal distance to removal (day)
Temporal distance to control event (day)
$\lambda_{obs\times10^2}$
$\lambda_{corr\times10^2}$

### Figures S2 and S3

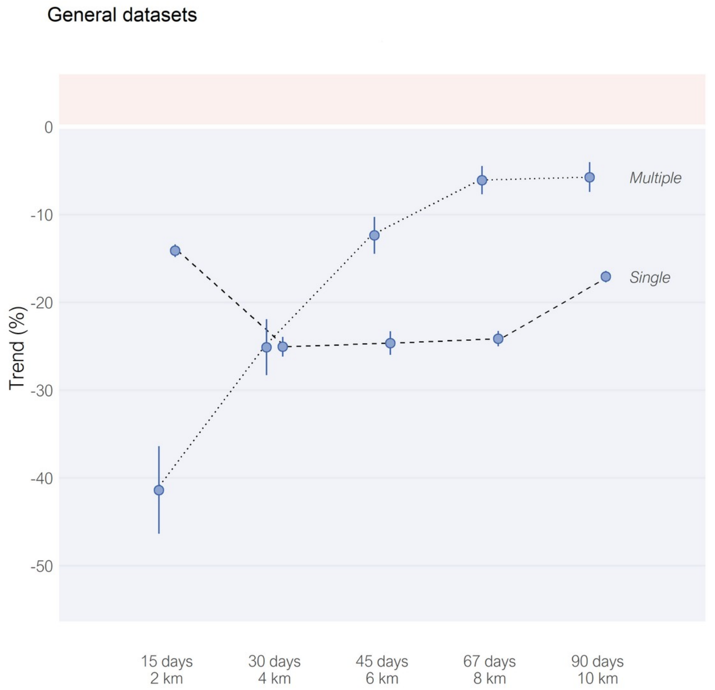


*Figure S2. Observed trends of attack intensities after the applications of single removals and multiple removals, at five nested scales. Points are mean estimates, alongside the confidence intervals (range 83.4%–90.0%). All trends are significant. Trends were not corrected for livestock presence.*

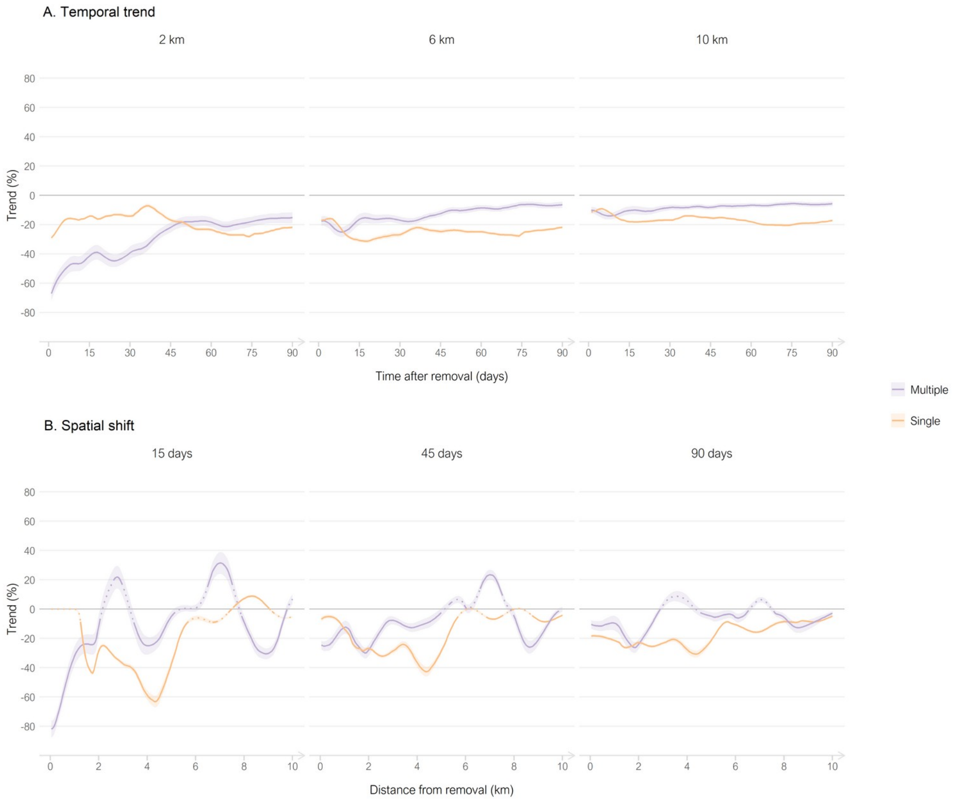


*Figure S3. Observed trends of attack intensities after the applications of single removals (orange) and of multiple removals (purple), alongside continuous temporal (A) and spatial (B) scales, for three fixed distances and periods, respectively. Lines are mean estimates, ribbons are the confidence intervals (range 83.4–95.0%). Dotted lines are not significant trends. Trends were not corrected for livestock presence. Note that temporal trends are calculated as nested periods (e.g. from 0 to x days, over 0–2 km for first panel), while spatial shifts are trends calculated for specific distances (e.g. at the $x^{th}$ km, over 15 days for first panel).*

### Figure S4

#### Administrative type (corrected)

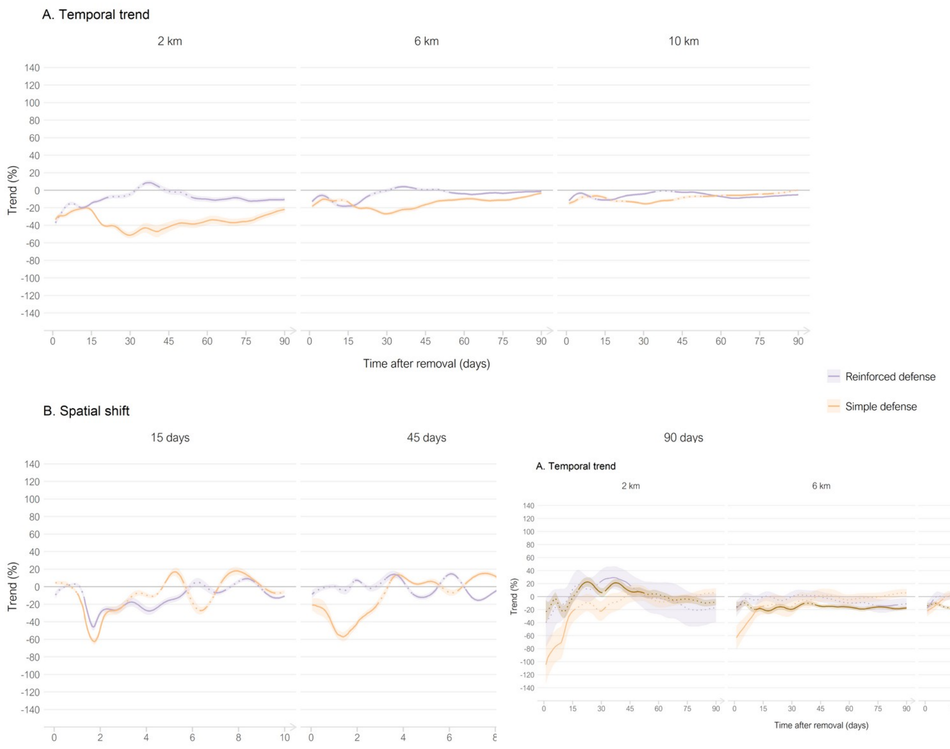


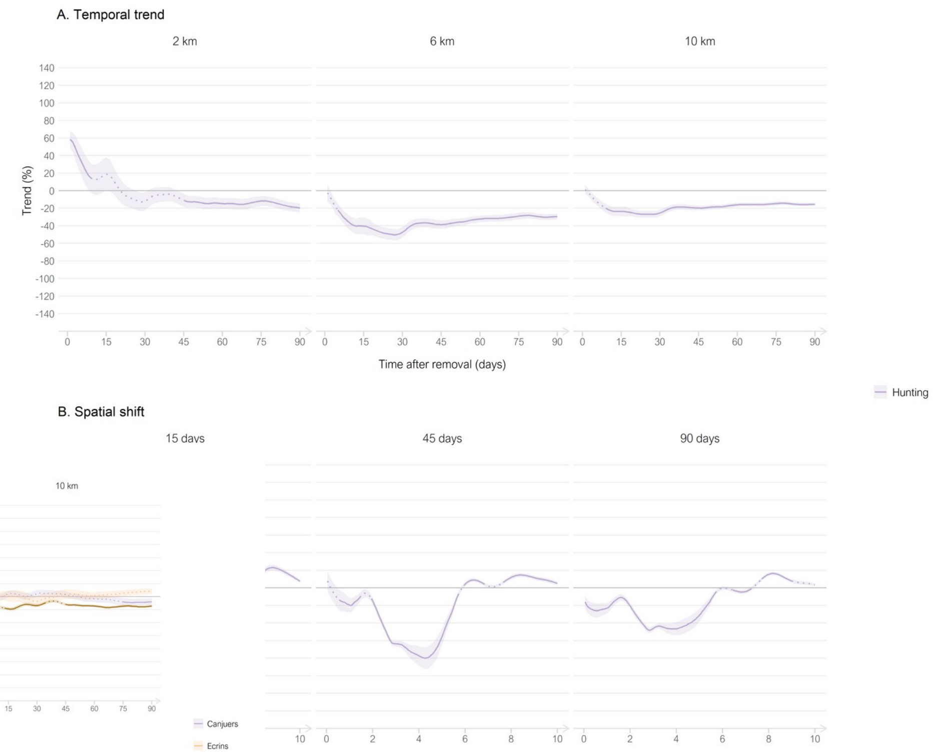


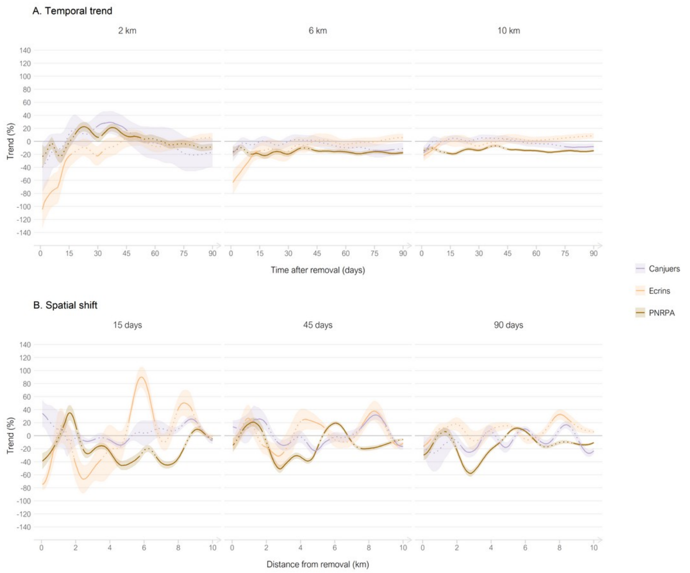


*Figure S4. Corrected trenc [illegible] 18 subsets, alongside continuous temporal (A) and spatial ( [illegible] mean estimates, ribbons are the confidence intervals (range [illegible] ed for livestock presence. Note that temporal trends are calcu [illegible] nel), while spatial shifts are trends calculated for specific dista [illegible]*

#### Geographic zone (corrected)

**Season (corrected)**

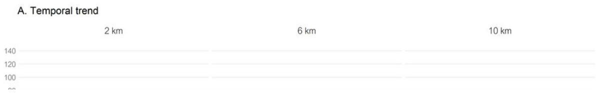

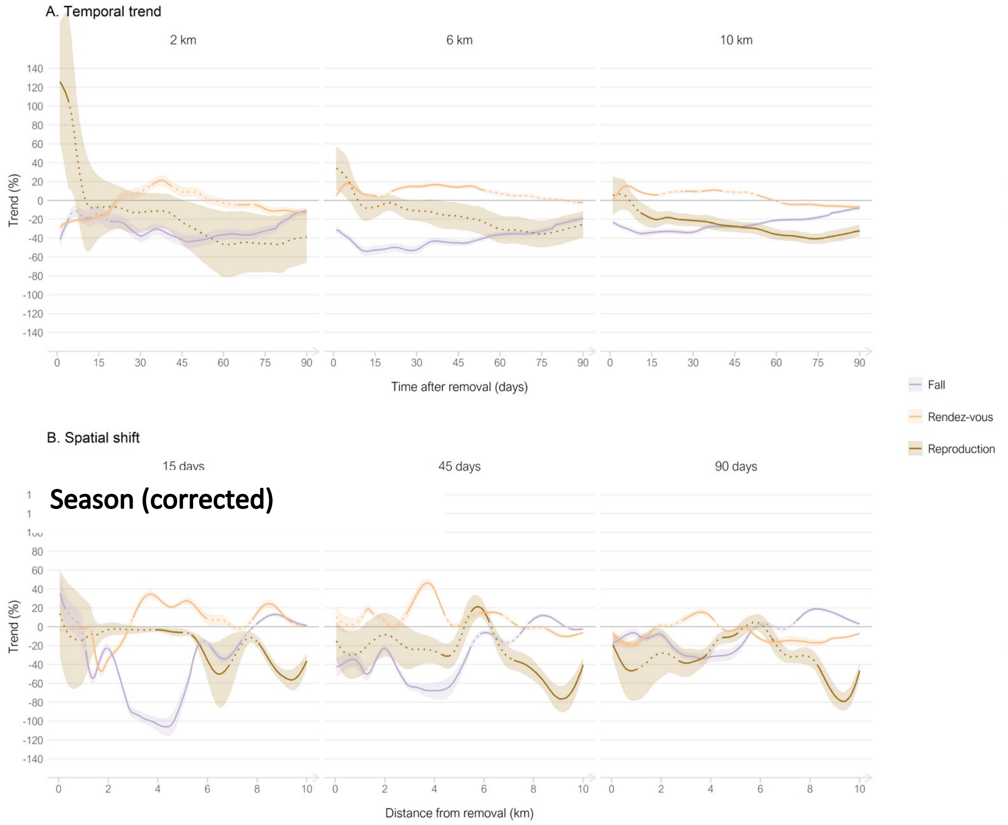


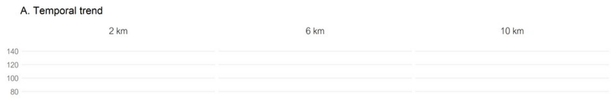

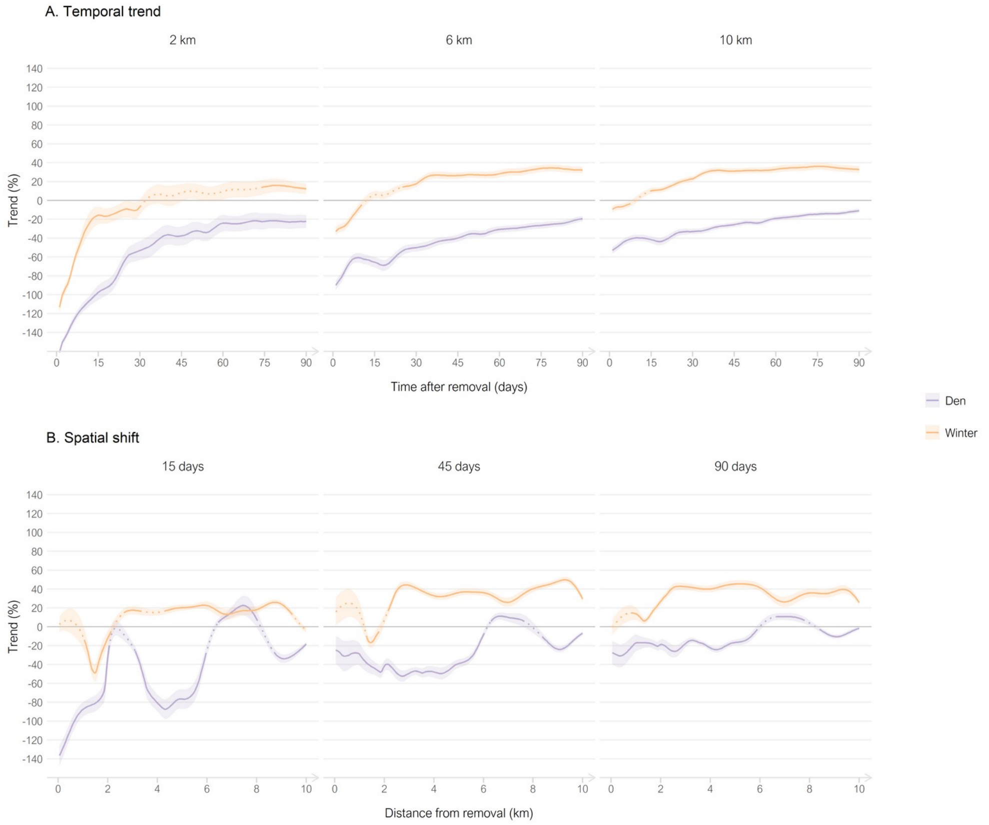

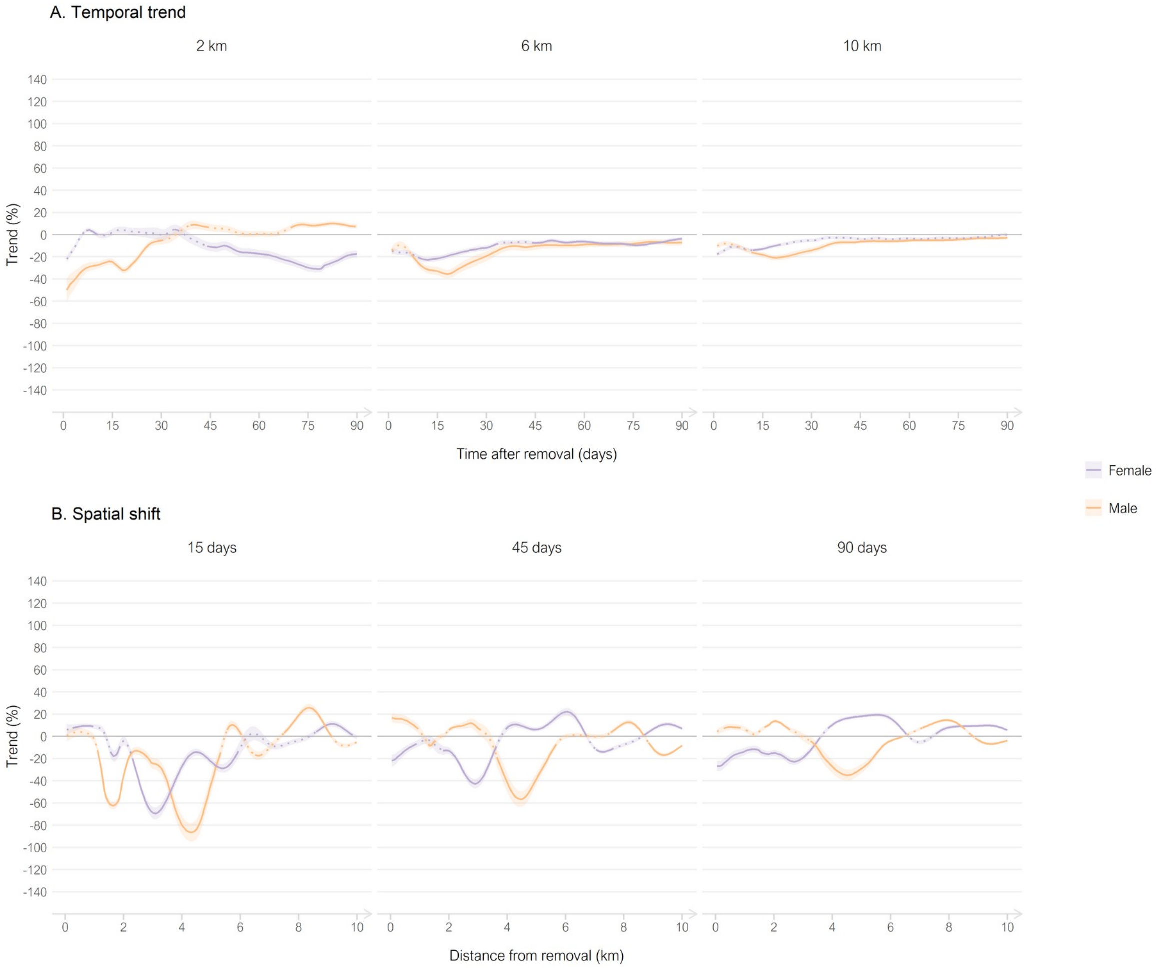


**Sex of the killed wolf (corrected)**

#### Figure S5

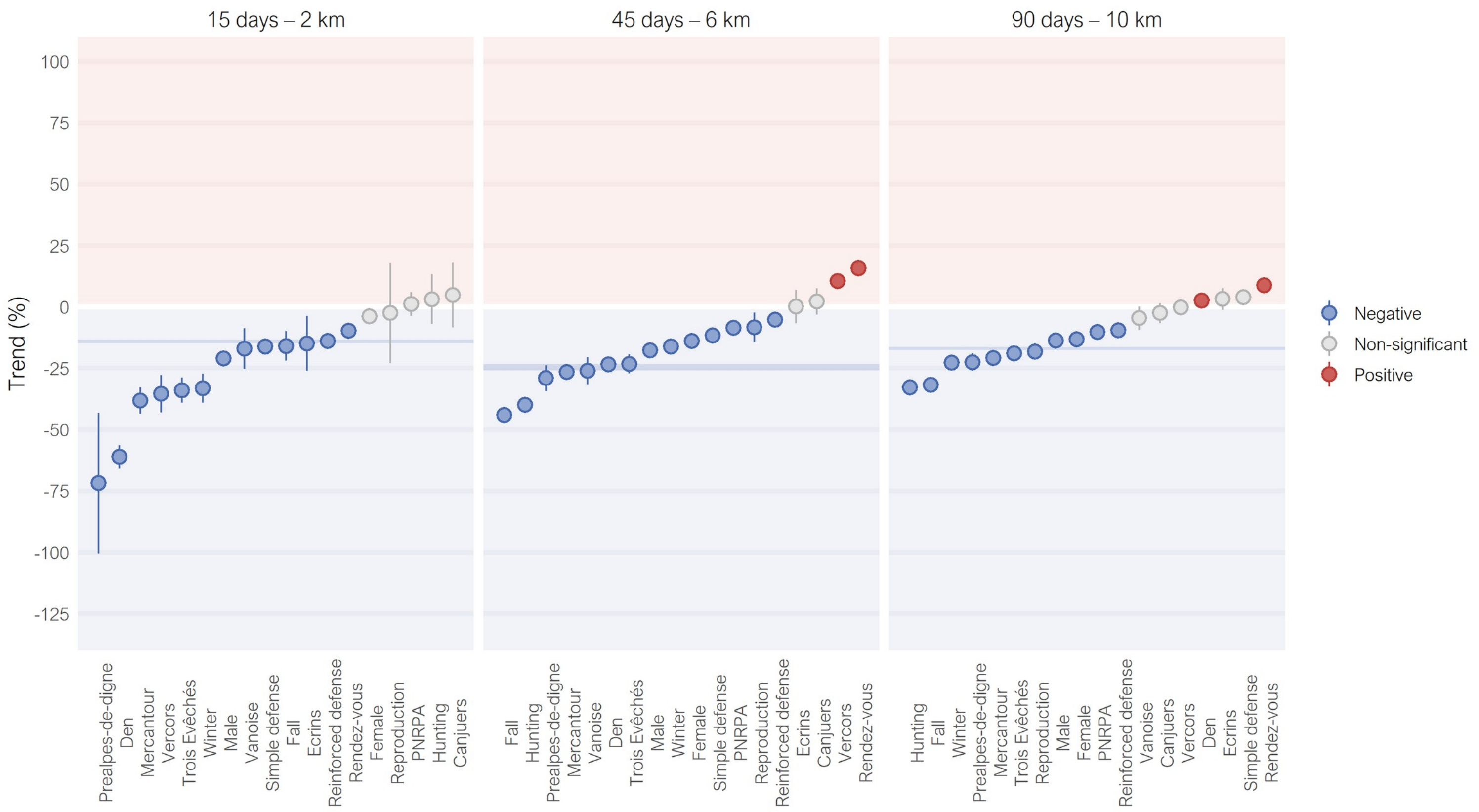


*Figure S5. Distributions of the observed trends of attack intensities for the 18 subsets of single removals at three nested scales. Points are mean estimates, alongside the confidence intervals (range 83.4%–87.9%). Trends in grey are not significant. Trends were not corrected for livestock presence. Confidence intervals of the entire dataset of single removals are in background in dark blue.*

#### Figure S6

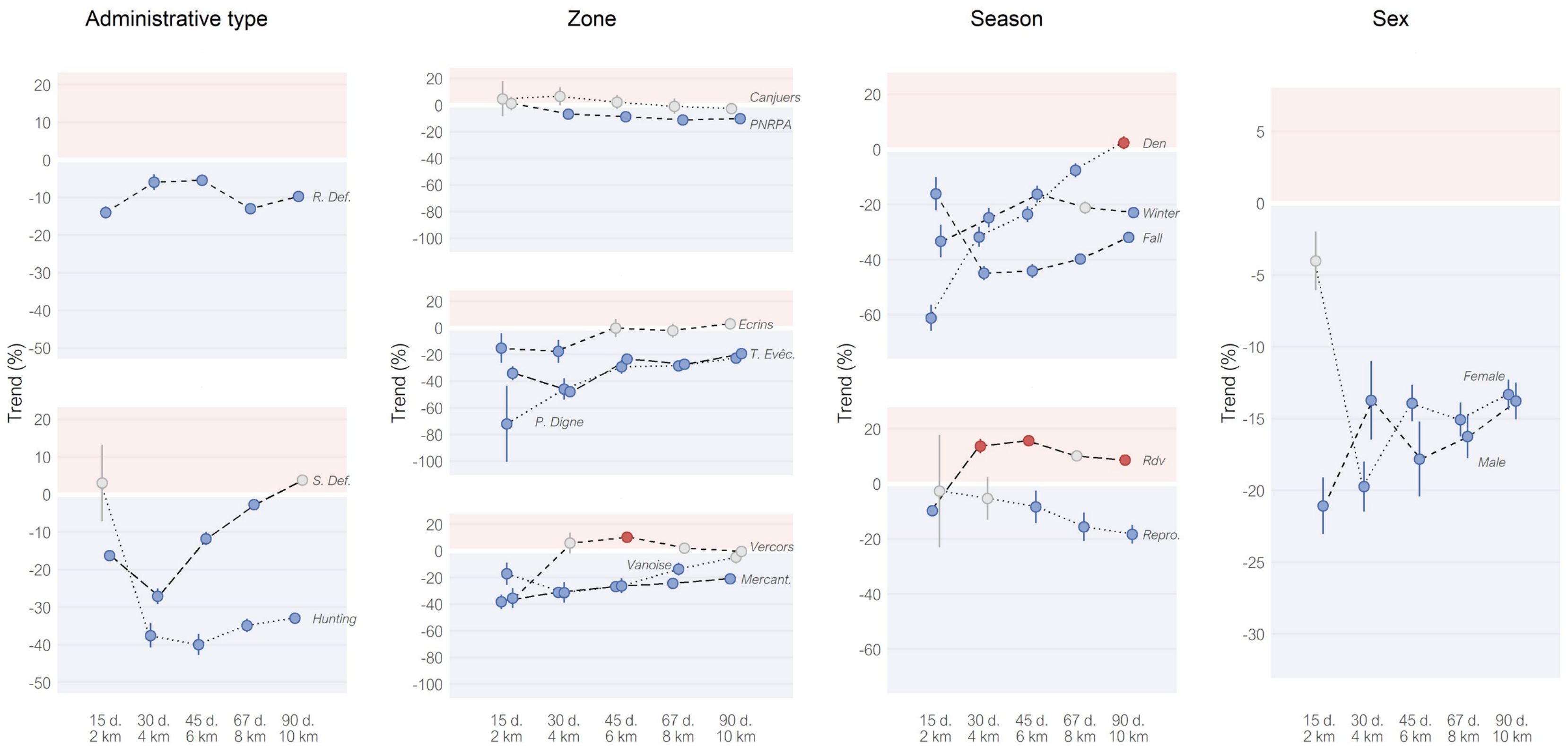


*Figure S6. Observed trends of attack intensities after the applications of single removals, divided into 18 subsets, at five nested scales. Points are mean estimates, alongside the confidence intervals (range 83.4%–87.9%). Trends in grey are not significant. Trends were not corrected for livestock presence. 'R. Def' and 'S. Def' for Reinforced and Simple defense subsets; 'T. Evêc.', 'P. Digne' and 'Mercant.' for Trois Evêchés, Préalpes de Digne and Mercantour subsets; 'Rdv' and 'Repro.' for Rendez-vous and Reproduction subsets.*

### Figure S7

#### Administrative type (not corrected)

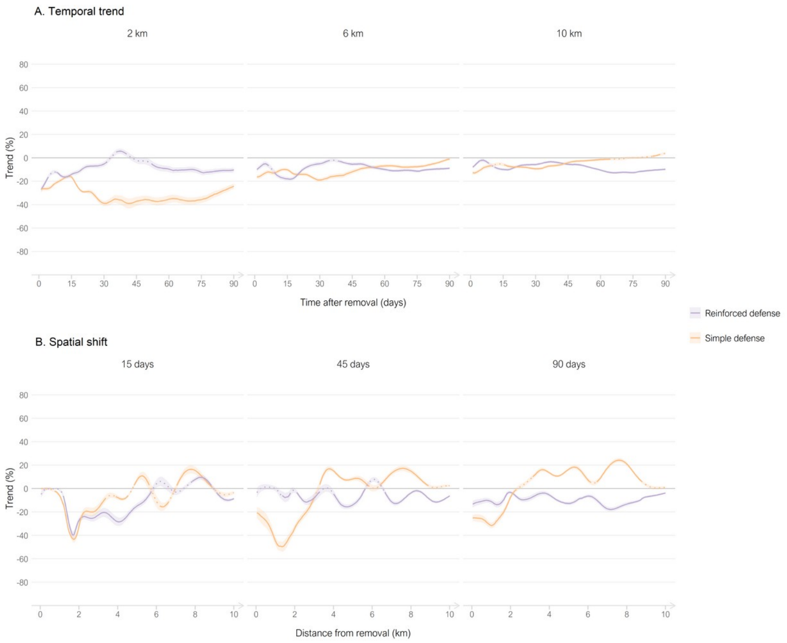


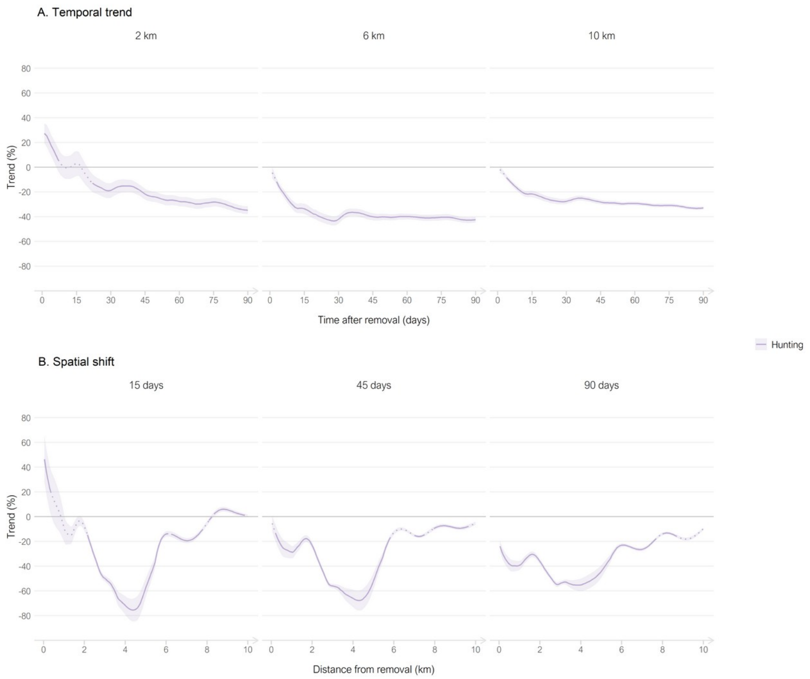


*Figure S4. Observed trends of attack intensities after the applications of single removals, for the 18 subsets, alongside continuous temporal (A) and spatial (B) scales, for three fixed distances and periods, respectively. Lines are mean estimates, ribbons are the confidence intervals (range 83.4–95.0%). Dotted lines are not significant trends. Trends were not corrected for livestock presence. Note that temporal trends are calculated as nested periods (e.g. from 0 to x days, over 0–2 km for first panel), while spatial shifts are trends calculated for specific distances (e.g. at the xth km, over 15 days for first panel).*

### Geographic zone (not corrected)

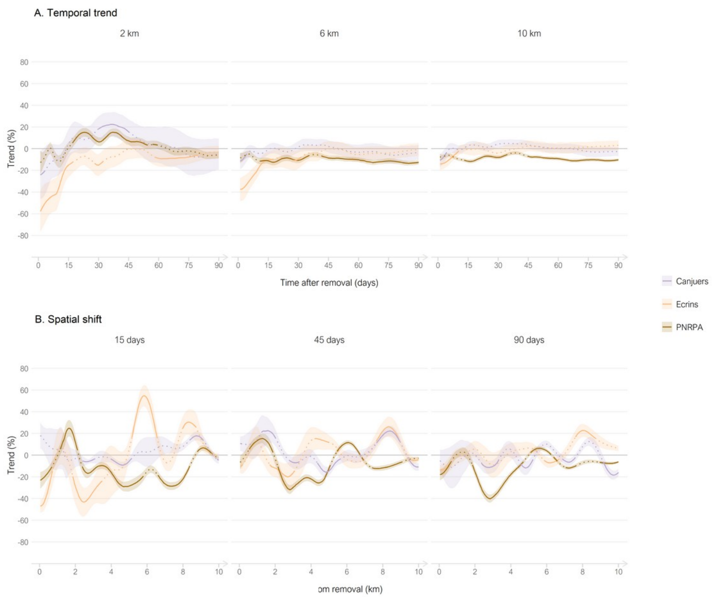


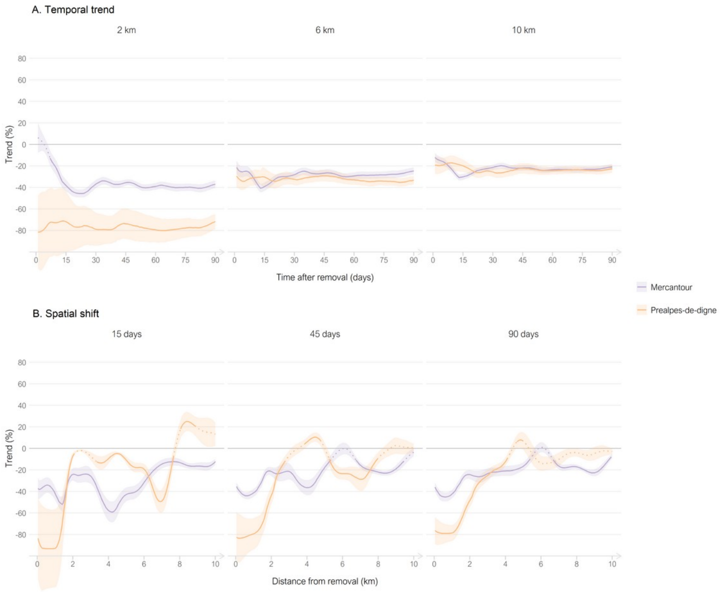


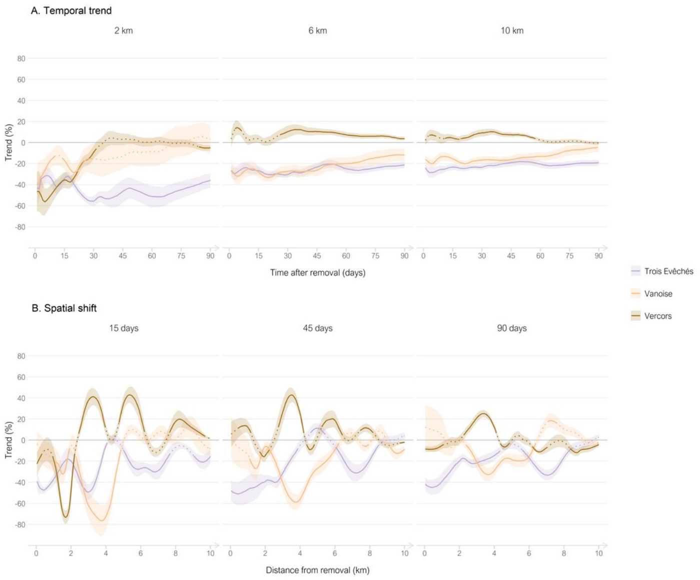

#### Season (not corrected)

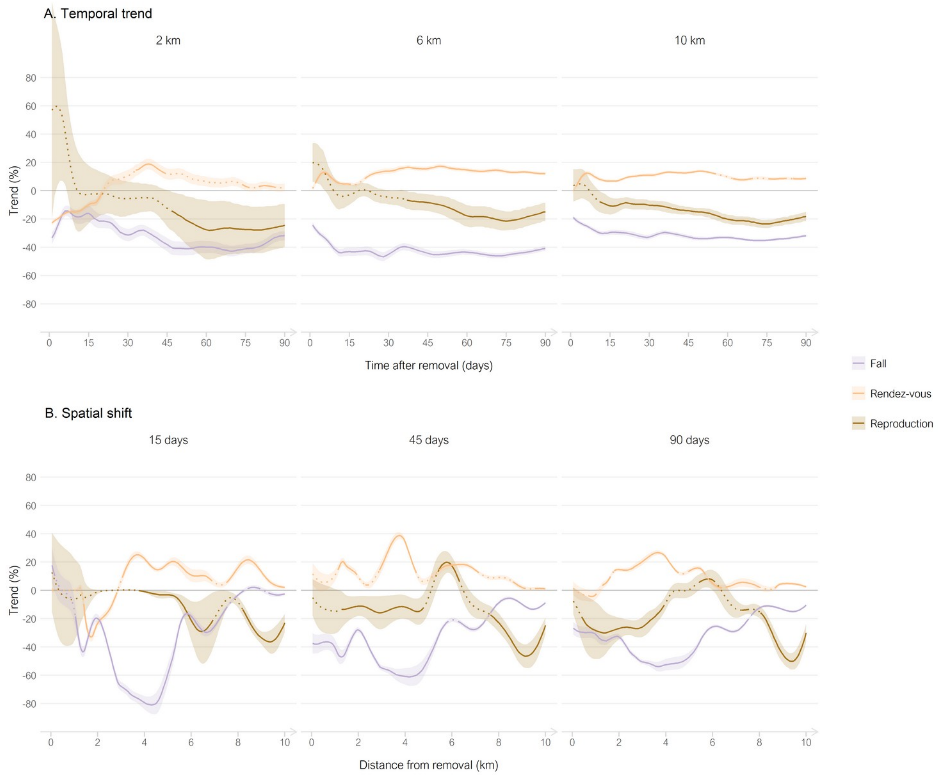


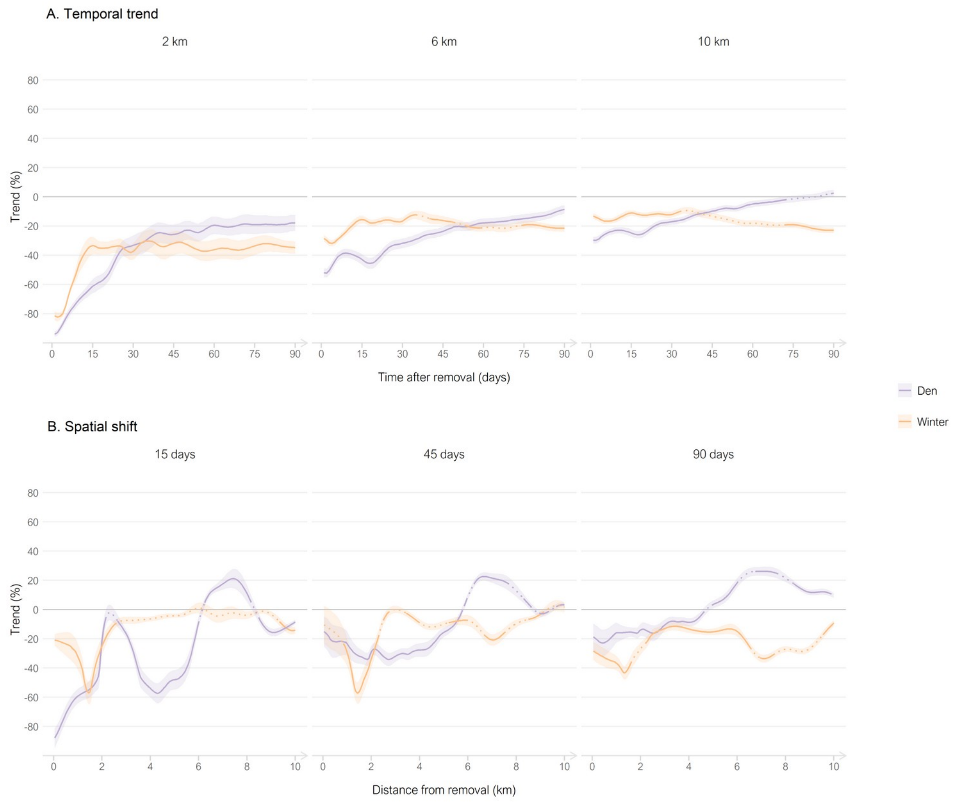

#### Sex of the killed wolf (not corrected)

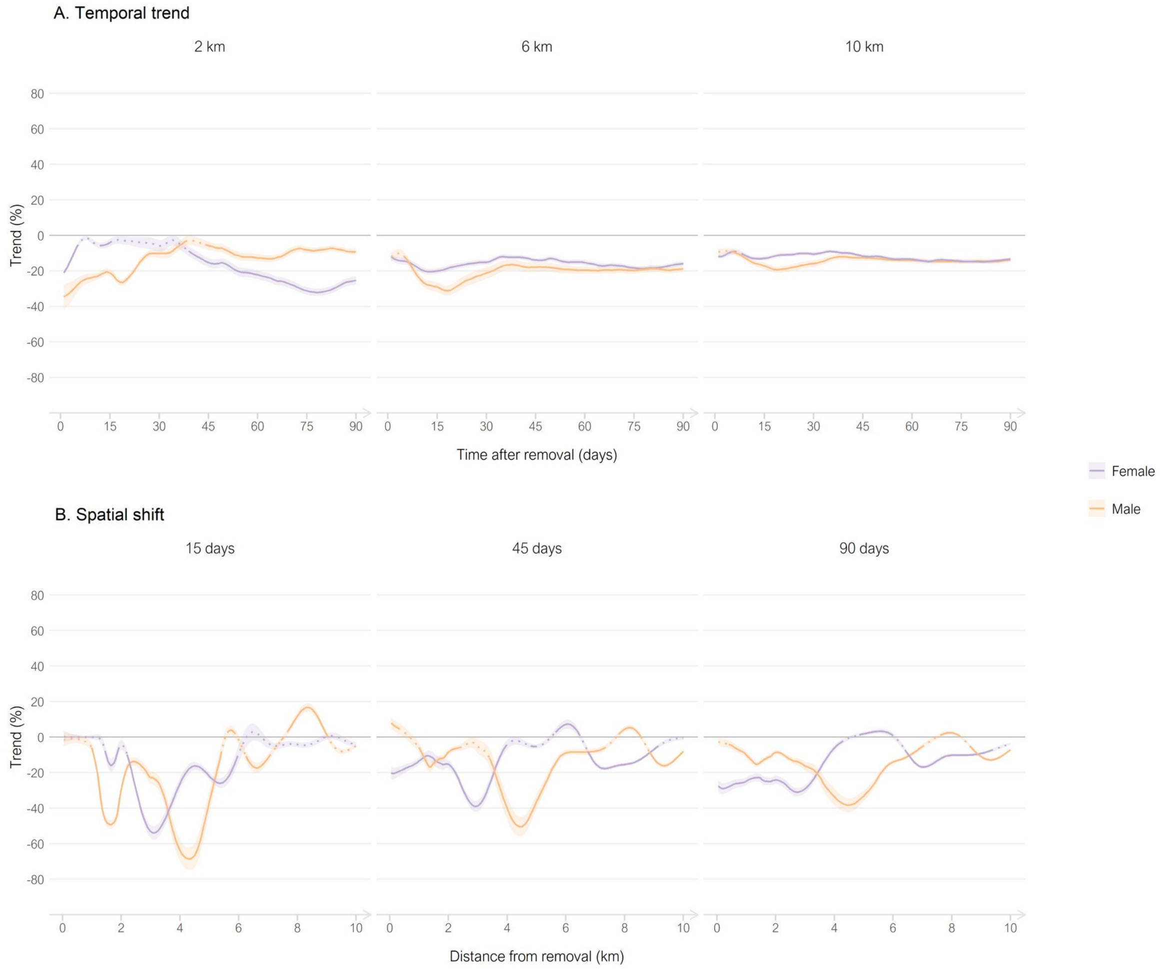

### Figure S8

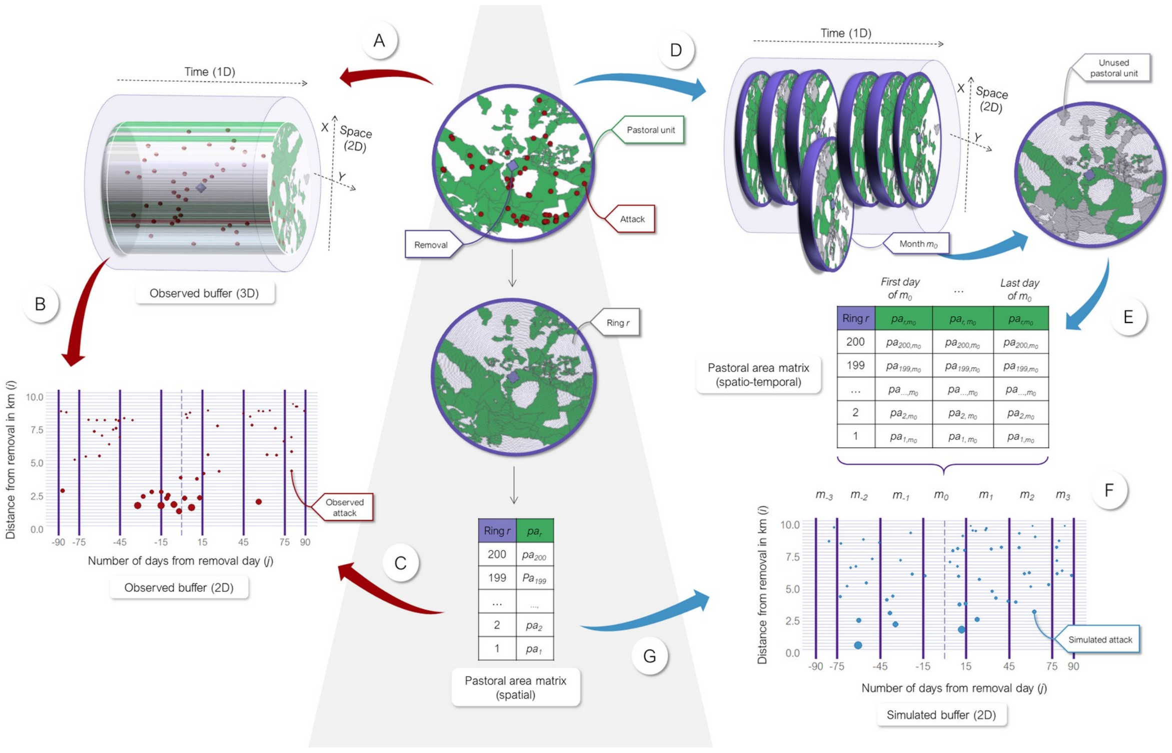


*Figure S8. Example of data transformations of observed attacks (red arrows) and simulated attacks (blue arrows) for one lethal removal occurring on the 15th of August 2017.* ***A.*** *Attribution of the observed 3D-buffer to the removal (purple point) of 10 km radius and 90 days around the removal. Attacks in the buffer have 3 coordinates: 1 temporal (day), and 2 spatial (x,y).* ***B.*** *Reduction of the spatial coordinates of the observed attacks, through the Euclidian distance to the removal (spatial coordinate i, time coordinate j).* ***C.*** *Spatial correction $1/pa_r$ showed as the size of the observed attacks. The pastoral area $pa_r$ is calculated for each of the 200 rings r of the spatial buffer.* ***D.*** *The seven monthly slices m of the observed 3D-buffer. Months could be complete (from $m_{-2}$ to $m_2$) or partial ($m_{-3}$ and $m_3$, here only 15 days each).* ***E.*** *The pastoral area $pa_{r,m0}$ is calculated for each of the 200 rings r of $m_0$. The unique column of $pa_{r,m0}$ is duplicated into the number of days of $m_0$. Note that the operation is repeated for each m, and results in a final matrix of 200 x 180 dimensions (not showed here).* ***F.*** *One simulation of coordinates i and j of attacks based on the final matrix of $pa_{r,m}$. The number of simulated attacks (blue points) of the simulated 2D-buffer is the same as the number of observed attacks (red points) of the observed buffer.* ***G.*** *Spatial correction $1/pa_r$ showed as the size of the simulated attacks, with the same process as in step C.*

#### Figure S9

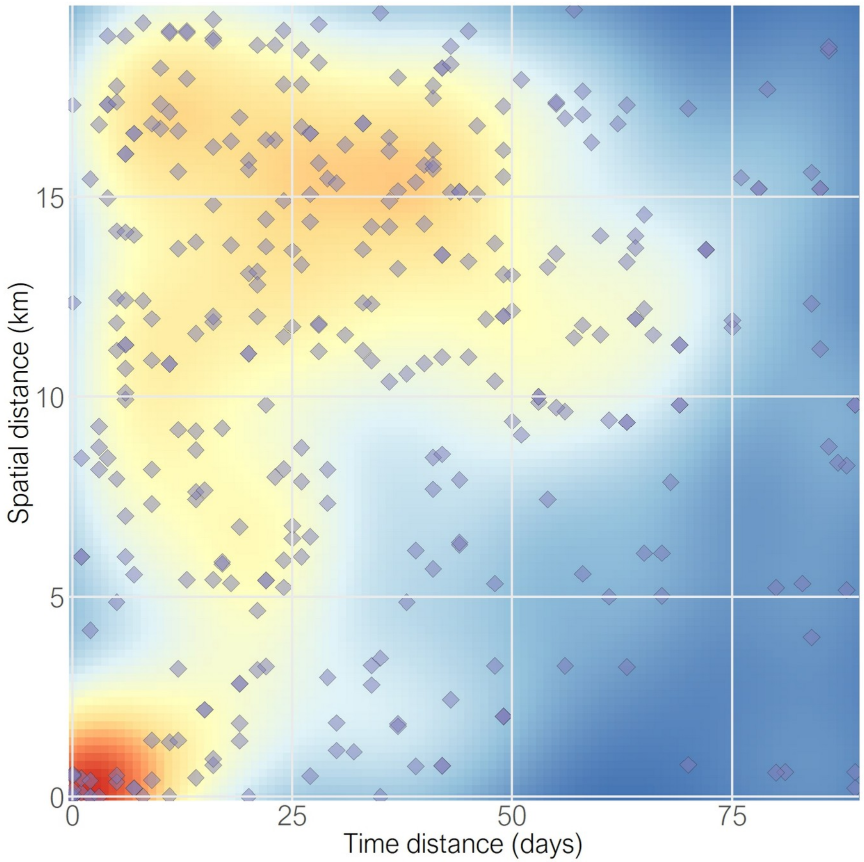


*Figure S9. Spatial and temporal distances between the 352 pairs of lethal removals whose buffers overlapped.*
