## Supplementary tables for "Spatio-temporal dynamics of attacks around deaths of wolves: A statistical assessment of lethal control efficiency in France"

*Table S1. Mean and confidence intervals of the trends of observed $Y_{\lambda_{obs}}$ and corrected $Y_{\lambda_{corr}}$ attack intensities for the lethal removals, calculated over three nested scales: 1 (0-2 km, ± 15 days), 2 (0-6 km, ± 45 days) and 3 (0-10 km, ± 90 days). Column "Total" indicates the number of removals of each dataset (single or multiple removals) or subset of single removals (for the other rows). Gradient of colors ranges from dark blue for -100 to dark red for 100, while the middle color in white corresponds to 0. Confidence intervals were adapted to each dataset or subsets (range 83.4–90.0%), following Knol et al., 2011. Significance of trends is not showed.*

| | | | Minimum CI | | | | | | Mean | | | | | | Maximum CI | | | | | |
|---|---|---|---|---|---|---|---|---|---|---|---|---|---|---|---|---|---|---|---|---|
| | | | Scale 1 | | Scale 2 | | Scale 3 | | Scale 1 | | Scale 2 | | Scale 3 | | Scale 1 | | Scale 2 | | Scale 3 | |
| | Subset | Total | $Y_{\lambda_{obs},1}$ | $Y_{\lambda_{corr},1}$ | $Y_{\lambda_{obs},2}$ | $Y_{\lambda_{corr},2}$ | $Y_{\lambda_{obs},3}$ | $Y_{\lambda_{corr},3}$ | $Y_{\lambda_{obs},1}$ | $Y_{\lambda_{corr},1}$ | $Y_{\lambda_{obs},2}$ | $Y_{\lambda_{corr},2}$ | $Y_{\lambda_{obs},3}$ | $Y_{\lambda_{corr},3}$ | $Y_{\lambda_{obs},1}$ | $Y_{\lambda_{corr},1}$ | $Y_{\lambda_{obs},2}$ | $Y_{\lambda_{corr},2}$ | $Y_{\lambda_{obs},3}$ | $Y_{\lambda_{corr},3}$ |
| | Single removal | 188 | -15 | -12 | -26 | -20 | -18 | -7 | -14 | -11 | -25 | -18 | -17 | -6 | -13 | -10 | -23 | -16 | -16 | -5 |
| | Multiple removal | 35 | -46 | -93 | -14 | -30 | -7 | -15 | -41 | -85 | -12 | -27 | -6 | -13 | -36 | -77 | -10 | -24 | -4 | -11 |
| Admin. type | *Reinforced defense* | 71 | -16 | -17 | -7 | -1 | -11 | -6 | -14 | -15 | -5 | 1 | -10 | -5 | -12 | -13 | -4 | 2 | -9 | -4 |
| | *Simple defense* | 65 | -18 | -25 | -14 | -18 | 3 | -1 | -16 | -23 | -12 | -16 | 4 | 0 | -15 | -20 | -10 | -14 | 5 | 2 |
| | *Hunting* | 52 | -7 | 1 | -43 | -43 | -34 | -17 | 3 | 19 | -40 | -39 | -33 | -15 | 13 | 38 | -37 | -34 | -32 | -14 |
| Geographic zone | *PNRPA* | 33 | -4 | -9 | -11 | -18 | -12 | -16 | 1 | -1 | -9 | -15 | -10 | -14 | 6 | 7 | -6 | -11 | -9 | -13 |
| | *Mercantour* | 23 | -44 | -62 | -30 | -27 | -23 | -12 | -38 | -53 | -27 | -22 | -21 | -10 | -33 | -44 | -23 | -17 | -18 | -7 |
| | *Trois Evêchés* | 18 | -39 | -37 | -27 | -8 | -22 | 1 | -34 | -30 | -23 | -3 | -19 | 6 | -29 | -23 | -19 | 2 | -16 | 11 |
| | *Vercors* | 18 | -43 | -49 | 7 | 25 | -2 | 16 | -36 | -38 | 10 | 30 | 0 | 19 | -28 | -28 | 14 | 35 | 1 | 22 |
| | *Vanoise* | 15 | -25 | -39 | -32 | -42 | -9 | -15 | -17 | -29 | -26 | -38 | -5 | -11 | -9 | -18 | -21 | -34 | 0 | -6 |
| | *Ecrins* | 12 | -26 | -40 | -7 | -8 | -1 | 5 | -15 | -22 | 0 | 2 | 3 | 9 | -4 | -4 | 7 | 12 | 8 | 13 |
| | *Préalpes de Digne* | 12 | -100 | -81 | -34 | -14 | -26 | 1 | -72 | -62 | -29 | -2 | -23 | 20 | -43 | -42 | -24 | 10 | -19 | 39 |
| | *Canjuers* | 11 | -9 | -10 | -3 | -8 | -7 | -13 | 5 | 11 | 2 | 0 | -3 | -8 | 18 | 32 | 8 | 8 | 1 | -3 |
| Season | *Fall* | 62 | -22 | -24 | -47 | -48 | -33 | -10 | -16 | -14 | -44 | -45 | -32 | -8 | -10 | -4 | -42 | -42 | -31 | -6 |
| | *Rendez-vous* | 56 | -11 | -19 | 14 | 12 | 8 | -9 | -10 | -16 | 16 | 14 | 9 | -7 | -8 | -14 | 17 | 16 | 10 | -5 |
| | *Denning* | 33 | -66 | -105 | -26 | -44 | 0 | -13 | -61 | -97 | -24 | -40 | 2 | -11 | -56 | -90 | -21 | -36 | 5 | -9 |
| | *Winter* | 29 | -39 | -25 | -19 | 23 | -25 | 29 | -33 | -16 | -16 | 27 | -23 | 33 | -27 | -7 | -13 | 30 | -21 | 36 |
| | *Reproduction* | 8 | -23 | -40 | -14 | -31 | -22 | -39 | -3 | -7 | -8 | -17 | -18 | -32 | 18 | 26 | -2 | -3 | -15 | -26 |
| Sex | *Male* | 80 | -23 | -28 | -20 | -13 | -15 | -4 | -21 | -25 | -18 | -10 | -14 | -3 | -19 | -21 | -15 | -7 | -12 | -1 |
| | *Female* | 74 | -6 | -1 | -15 | -9 | -14 | -1 | -4 | 2 | -14 | -8 | -13 | 0 | -2 | 5 | -13 | -6 | -12 | 1 |

*Table S2. Mean and confidence intervals of the trends of observed $Y_{\lambda_{obs}}$ and corrected $Y_{\lambda_{corr}}$ attack intensities for the 100 sets of control events, calculated over three nested scales: 1 (0-2 km, ± 15 days), 2 (0-6 km, ± 45 days) and 3 (0-10 km, ± 90 days). Column "Total" indicates the number of removals of each dataset (single or multiple removals) or subset of single removals (for the other rows). Gradient of colors ranges from dark blue for -100 to dark red for 100, while the middle color in white corresponds to 0. Confidence intervals were adapted to each dataset or subsets (range 83.4–90.0%), following Knol et al., 2011.*

| | | | Minimum CI | | | | | | Mean | | | | | | Maximum CI | | | | | |
|---|---|---|---|---|---|---|---|---|---|---|---|---|---|---|---|---|---|---|---|---|
| | | | Scale 1 | | Scale 2 | | Scale 3 | | Scale 1 | | Scale 2 | | Scale 3 | | Scale 1 | | Scale 2 | | Scale 3 | |
| | Subset | Total | $Y_{\lambda_{obs},1}$ | $Y_{\lambda_{corr},1}$ | $Y_{\lambda_{obs},2}$ | $Y_{\lambda_{corr},2}$ | $Y_{\lambda_{obs},3}$ | $Y_{\lambda_{corr},3}$ | $Y_{\lambda_{obs},1}$ | $Y_{\lambda_{corr},1}$ | $Y_{\lambda_{obs},2}$ | $Y_{\lambda_{corr},2}$ | $Y_{\lambda_{obs},3}$ | $Y_{\lambda_{corr},3}$ | $Y_{\lambda_{obs},1}$ | $Y_{\lambda_{corr},1}$ | $Y_{\lambda_{obs},2}$ | $Y_{\lambda_{corr},2}$ | $Y_{\lambda_{obs},3}$ | $Y_{\lambda_{corr},3}$ |
| | Single removal | 188 | -3 | -5 | -4 | -5 | -3 | 0 | 0 | 1 | -3 | -4 | -2 | 1 | 4 | 6 | -2 | -2 | -2 | 2 |
| | Multiple removal | 35 | 3 | -3 | 5 | 0 | 4 | -1 | 6 | 2 | 6 | 1 | 4 | 1 | 9 | 7 | 7 | 3 | 5 | 2 |
| Admin. type | *Reinforced defense* | 71 | -4 | -7 | 0 | -1 | 2 | 2 | -1 | -1 | 1 | 1 | 2 | 3 | 3 | 5 | 2 | 2 | 3 | 4 |
| | *Simple defense* | 65 | -1 | -4 | -2 | -8 | 4 | -1 | 3 | 1 | -1 | -7 | 4 | 0 | 6 | 6 | 0 | -5 | 5 | 1 |
| | *Hunting* | 52 | -7 | -6 | -12 | -3 | -17 | 0 | -3 | 0 | -11 | -1 | -16 | 2 | 0 | 6 | -9 | 1 | -15 | 4 |
| Geographic zone | *PNRPA* | 33 | -7 | -12 | -5 | -8 | -5 | -10 | -5 | -9 | -3 | -6 | -4 | -8 | -2 | -5 | -2 | -5 | -3 | -7 |
| | *Mercantour* | 23 | -1 | -2 | -3 | -2 | -5 | 1 | 2 | 3 | -2 | 0 | -4 | 2 | 5 | 8 | -1 | 2 | -3 | 3 |
| | *Trois Evêchés* | 18 | -7 | -5 | -3 | 7 | -5 | 10 | -3 | 1 | -2 | 9 | -4 | 11 | 0 | 7 | 0 | 11 | -3 | 12 |
| | *Vercors* | 18 | -5 | -10 | -1 | 1 | -1 | 3 | -1 | -4 | 0 | 3 | 0 | 4 | 2 | 1 | 2 | 5 | 2 | 5 |
| | *Vanoise* | 15 | -5 | -11 | 0 | -5 | 6 | 2 | -2 | -6 | 1 | -3 | 7 | 3 | 1 | 0 | 2 | -1 | 7 | 4 |
| | *Ecrins* | 12 | -3 | -4 | 1 | -2 | 5 | 8 | 1 | 2 | 2 | 1 | 6 | 9 | 5 | 8 | 4 | 3 | 7 | 10 |
| | *Préalpes de Digne* | 12 | -1 | 5 | -4 | 3 | -5 | 4 | 2 | 9 | -2 | 5 | -4 | 6 | 6 | 14 | -1 | 8 | -3 | 8 |
| | *Canjuers* | 11 | 2 | -5 | 4 | 0 | 4 | -1 | 6 | 0 | 6 | 3 | 5 | 1 | 10 | 6 | 7 | 5 | 6 | 2 |
| Season | *Fall* | 62 | -9 | -12 | -18 | -13 | -17 | 0 | -6 | -6 | -16 | -11 | -17 | 1 | -3 | -1 | -15 | -9 | -16 | 3 |
| | *Rendez-vous* | 56 | 2 | 1 | 7 | 5 | 10 | 4 | 5 | 7 | 8 | 7 | 11 | 5 | 8 | 12 | 9 | 9 | 11 | 6 |
| | *Denning* | 33 | -1 | -24 | 10 | -7 | 20 | 1 | 1 | -19 | 11 | -4 | 21 | 3 | 4 | -14 | 12 | -2 | 21 | 5 |
| | *Winter* | 29 | -7 | 6 | -13 | 7 | -28 | -2 | -4 | 11 | -12 | 8 | -27 | -1 | -1 | 17 | -11 | 10 | -26 | 1 |
| | *Reproduction* | 8 | 3 | -4 | -1 | -15 | -7 | -22 | 6 | 1 | 0 | -13 | -6 | -20 | 8 | 6 | 2 | -11 | -5 | -19 |
| Sex | *Male* | 80 | -3 | -5 | -4 | -4 | -3 | 2 | 0 | 1 | -3 | -2 | -2 | 3 | 4 | 6 | -2 | -1 | -2 | 4 |
| | *Female* | 74 | -3 | -6 | -4 | -5 | -3 | -1 | 0 | -1 | -3 | -3 | -3 | 0 | 4 | 4 | -2 | -2 | -2 | 1 |
